# Single-Cell Profiling Reveals Altered Endometrial Cellular Features Across the Menstrual Cycle in Endometriosis Patients

**DOI:** 10.1101/2025.07.21.666016

**Authors:** Ana Almonte-Loya, Wanxin Wang, Sahar Houshdaran, Xinyu Tang, Emily Flynn, Binya Liu, Divyashree Kushnoor, Camran Nezhat, Felipe Vilella, Carlos Simon, Kim Chi Vo, Amber Casillas, Umair Khan, Carlota Peredas, Tomiko T. Oskotsky, David Huang, Júlia Vallvé-Juanico, Juan Irwin, Alexis Combes, Marina Sirota, Gabriela Fragiadakis, Linda Giudice

## Abstract

Endometriosis is a chronic, estrogen-dependent condition affecting over 190 million women globally, characterized by the ectopic presence of endometrial-like tissue that leads to inflammation, pain, and infertility. Despite its prevalence, the pathogenesis of endometriosis remains poorly understood. Here, we present a comprehensive single-cell transcriptomic atlas comprising 228,000 cells derived from 43 eutopic endometrial biopsies from patients with endometriosis, fibroid controls and healthy controls, sampled across the menstrual cycle. This analysis reveals previously uncharacterized subpopulations of endometrial fibroblasts and epithelial cells undergoing epithelial–mesenchymal transition, alongside disrupted immune cell communication networks. Comparative gene expression profiling implicates oxidative stress, aberrant cell migration, and dysregulated apoptosis as central features of the disease state. These findings suggest that endometriosis alters eutopic endometrial homeostasis, with potential consequences for fertility, regeneration, and disease progression. Our dataset provides a valuable resource for biomarker discovery and identifies candidate therapeutic targets aimed at restoring endometrial function and alleviating symptoms in affected individuals.

## Introduction

Endometriosis is a common, estrogen-dependent inflammatory disease characterized by the ectopic growth of endometrial-like tissue outside the uterus. Most disease resides in the pelvia and is associated with inflammation, chronic pelvic pain, pain with intercourse and menses, and and infertility^1–5^. It affects around 10% of reproductive-age individuals with a uterus^2–4^, equating to approximately 190 million cases worldwide^2,6^. The economic burden in the U.S. alone exceeds $22 billion^7,8^. Despite decades of research and some advancements in exploring the disease’s pathobiology, effective individualized management remains challenging, primarily because of the heterogeneous nature of endometriosis phenotypes and symptoms, and involvement of systemic components^2,4^.

Although retrograde menstruation is a widely proposed origin of endometriosis, bone marrow stem cells may also contribute to its pathogenesis, and understanding eutopic (within the uterus) endometrial alterations in patients with disease remains limited, particularly in relation to tissue dyshomeostasis throughout the menstrual cycle and impact on fertility and pregnancy outcomes^1–4,9^. Since the endogenous ovarian steroids, estradiol and progesterone, are key regulators of endometrial gene expression^9–11^, investigating endometrial features across the menstrual cycle is key to understanding how aberrancies of signaling by these hormones drive endometrial dysfunction in endometriosis patients^4,9^.

Single-cell analysis has emerged as a powerful tool to decipher the complexities of diseases like endometriosis, offering detailed insights into the cellular heterogeneity of the eutopic and ectopic endometrium^9,12–17^. Recent studies utilizing these advanced methodologies have revealed diverse roles of immune cells, stromal fibroblasts, and epithelial cells in the eutopic endometrium proper and in relation to tissue dysfunction and pathogenesis and pathophysiology of endometriosis lesions. These studies have shown that eutopic endometrium in endometriosis patients exhibits epithelial abnormalities and dysregulated signaling pathways affecting tissue homeostasis. Moreover, analysis of menstrual endometrium, a “liquid biopsy” of late secretory phase endometrium, reveals aberrant progesterone signaling in stromal fibroblasts, which are essential for pregnancy establishment and success and endometrial regeneration^9,13^. These findings underscore the importance of cellular diversity and functional interactions in the eutopic endometrium of patients with endometriosis and highlight potential avenues for targeted therapeutic interventions.

In this study, we utilized single-cell RNA sequencing (scRNA-seq) to examine cellular and molecular changes as well as cell-cell communication patterns in the eutopic endometrium of patients with and without endometriosis across disease severities and menstrual cycle phases. By profiling the eutopic endometrium, we aimed to identify cycle phase-dependent alterations in key cell populations, including stromal fibroblasts, unciliated epithelial cells, EMT cells, and immune dysregulation in macrophages and uterine natural killer (uNK) cell types and subtypes. Through these analyses, we sought to map critical cell-cell signaling pathways involved in inflammation, apoptosis, and migration, offering deeper insights into the molecular mechanisms driving endometrial dysfunction in patients with endometriosis.

## Results

### Diverse cellular cartography of the eutopic endometrium

By using multiplexed single-cell RNA sequencing to mitigate batch effect and other technical artifacts, single-cell analyses identified 228,021 cells from the eutopic endometrium of 43 participants, including 25 with endometriosis, 8 with no known reproductive disorders (“pristine” controls), and 10 with uterine fibroids and no endometriosis (“fibroid” controls). Patient samples were stratified according to case/control, menstrual cycle phase and endometriosis severity^18^ (**Fig. 1A, Supplementary Table 1**). Following experimental processing, samples were processed in silico including quality filtering, integration, and batch correction, prior to downstream computational analysis (**Fig. 1B, Methods**).

**Figure 1:**
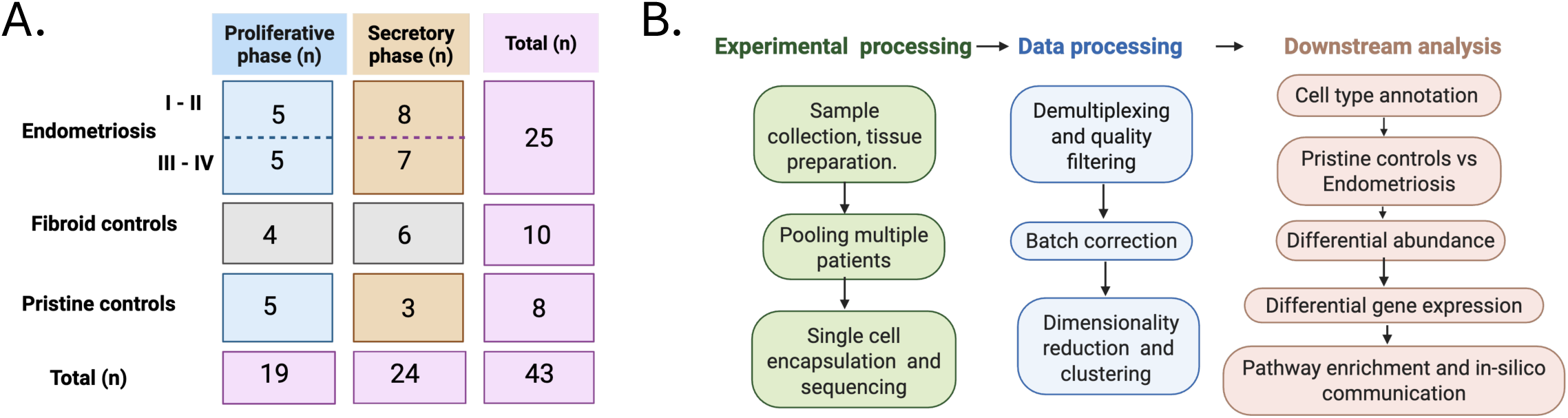
Data and preprocessing. A. Patient cohort and number of samples p srofiled. Classified by disease status and menstrual cycle phase. B. Experimental and bioinformatic processing workflow summary.

Using established cell markers and visualizing using Uniform Manifold Approximation and Projection (UMAP) visualization, we identified thirty-two clusters representing six major cell types: immune cells, stromal fibroblasts, ciliated epithelial cells, unciliated epithelial cells, endothelial cells, and vascular smooth muscle cells (VSM) (**Fig. 2A, SFig. 1A, STable 2**). We then explored the distribution of menstrual phases and disease states within these cell clusters. Coloring the UMAP based on these characteristics showed distinct regions for proliferative and secretory phases (**Fig. 2B and SFig. 1B**), as well as separations between endometriosis, fibroid control, and pristine control (**Fig. 2C and SFig. 1C**). Menstrual phase, disease status, and libraries were distributed across all cell types (**SFig. 1D-G**). Principal component analysis (PCA) of the cell composition profile per sample further confirmed changes in endometrial cellular composition driven by menstrual cycle phase and disease condition (**Fig. 2D,E**). Notably, the separation among conditions (endometriosis, fibroid controls, and pristine controls) was less pronounced than differences driven by cycle phase. Similar to the cellular composition analysis results, most cell types showed significant transcriptional differences between proliferative and secretory phases in fibroblast and unciliated epithelia populations (**SFig. 2-4, Supplementary Note, STable4)**. Interestingly, although endometrium from patients affected by fibroids exhibited distinct signals, their molecular signatures were more similar to endometriosis than to pristine controls in certain cell types (examples in **SFig.5, Supplementary Note below, STable5**). Thus, we focused downstream analyses on comparing eutopic endometrium signatures in endometriosis and pristine controls, stratified by menstrual cycle phase (**Fig. 2F**).

**Figure 2:**
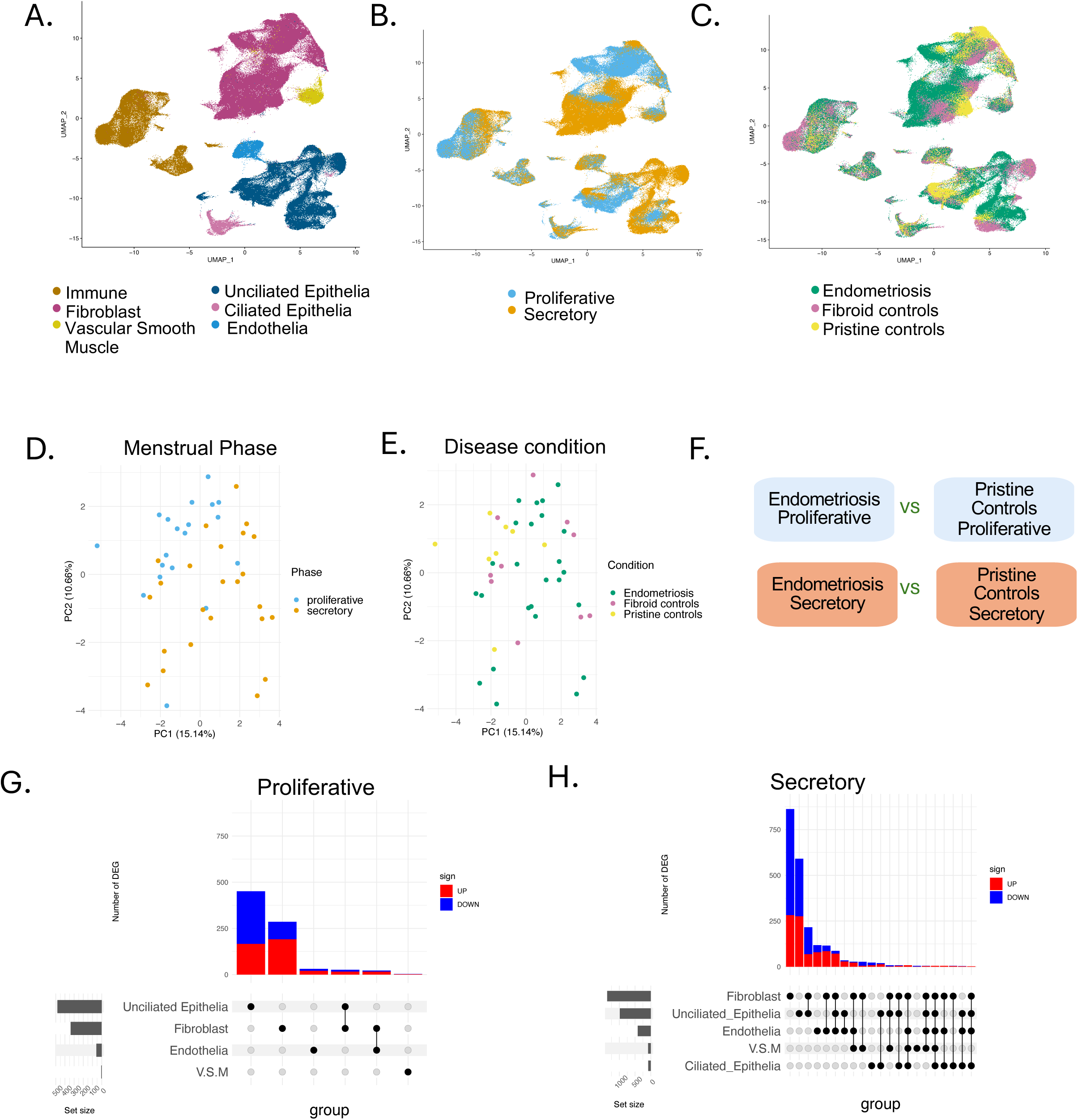
Single-cell analysis of endometrial tissue to examine the molecular differences in endometriosis patients across disease states and menstrual cycle phase. A. Global UMAP visualization of all sequenced cells, depicting six major broad cell types. B. UMAP plot colored by the menstrual cycle phase of the samples. C. UMAP plot displaying samples by disease status: endometriosis, pristine controls, and fibroid controls. D. PCA composition colored by menstrual cycle phase. E. PCA composition colored by disease status. F. Study design for differential gene expression analysis. Samples were stratified by phase to compare pristine controls with endometriosis samples. Differential expression was assessed with P<0.05 and log2 fold change (log2 FC) > 0.25. Analyses were performed using MAST, with p values adjusted. G. Differentially expressed genes in stratified proliferative samples across major cell types. Upregulated genes in endometriosis are shown in red (log2 FC > 0.25), while downregulated genes are shown in blue (log2 FC < -0.25). An UpSet plot illustrates the overlap of genes exhibiting expression changes across different cell types. H. Differentially expressed genes in stratified secretory samples across major cell types

### Differential gene expression across cell types in endometrial pathology

We first examined differentially expressed genes (DEGs) filtering by adj. p-val < 0.05 and |log2FC| > 0.25, between endometriosis and pristine controls in the five non-immune broad cell types (**STable 3**) (**SFig. 6**). In both the proliferative and secretory phrases, we saw the highest numbers of DEGs in stromal fibroblasts (n=338 and n=1384 respectively) followed by unciliated epithelial cells (n=482 and n=983) and endothelial cells (n=58, n=417). VSM and ciliated epithelial cells also showed DEGs, mainly in the secretory phase (n=5 and n=88 for VSM, n=4 and n=84 for CE) (**SFig. 6**).

We found that all non-immune cell types had more DEGs in the secretory phase than in the proliferative phase (**Fig. 2G, H**). Although most gene differences generally appeared to be phase-specific (**SFig. 7A-E.**), there was greater overlap of DEGs across cell types in the secretory phase compared to the proliferative phase (**Fig. 2G, H**).

These findings suggest that endometriosis exerts greater transcriptional effects in secretory phase endometrium, which may have significant implications for fertility and reproductive health. To investigate if the observed DEGs are isolated to specific endometriosis severity groups, we also performed similar comparisons stratified by endometriosis stage, which showed general concordance with the transcriptomic differences seen in the analyses without endometriosis severity stratification (**SFig8, Supplementary Notes, STable6,7**). Therefore, we continued with an analysis approach unstratified by endometriosis stage.

### Distinct biological process alterations between menstrual cycle phases in stromal fibroblasts in endometriosis cases

We decided to investigate the two main mesenchymal-derived cell populations, stromal fibroblast and VSM (**Fig. 3A**). Given the large number of significant DEGs in stromal fibroblasts in endometriosis compared to pristine controls, we first focused the on the signatures of this cell population. We found 204 overlapping DEGs between the proliferative phase and secretory phase, among which 156 genes consistently exhibited either upregulation or downregulation in both phases in patients with endometriosis compared to pristine controls (**Fig. 3B, SFig. 9A**).

**Figure 3:**
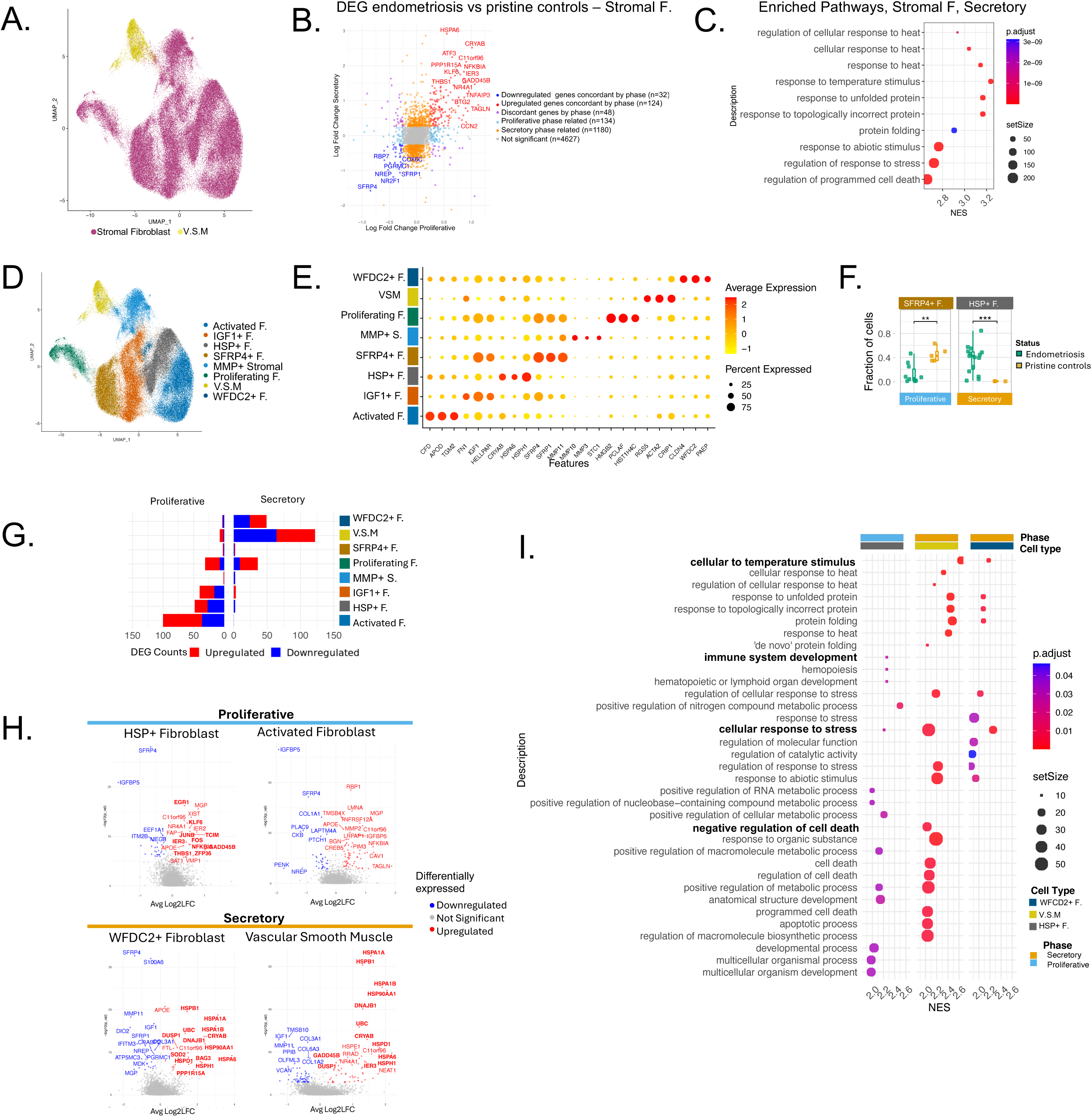
Phase-specific expression in stromal fibroblasts and vascular smooth muscle cells reveals stress, apoptosis, and immune response pathways in endometriosis patients. A. UMAP visualization of stromal cell subtypes, color-coded by major tissue type classes (as shown in Fig. 1A). B. Log2FoldChange of DEG in the secretory and proliferative phase, comparing pristine controls and endometriosis samples. Samples were stratified by menstrual phase. DGE was assessed using a threshold of p-adj.<0.05 and |log2FC| > 0.25. C. Upregulated pathways in all stromal populations in secretory phase. Showing Normalized Enrichment Score (NES), p-adj <0.05. D. UMAP visualization of 8 stromal cell subtypes, color-coded by subtype identity. E. Top marker genes for each of the 8 stromal cell subtypes. F. Boxplot showing differential abundance of stromal cell types across samples. Statistical significance was determined using a t-test and corrected p-values. G. Number of DEGs comparing control and endometriosis samples for each stromal subtype, stratified by menstrual phase. Red, upregulated in endometriosis, blue downregulated in endometriosis. H. Differentially regulated genes between pristine controls and endometriosis samples, stratified by menstrual phase. Bolded genes are involved in selected pathways of stress, apoptosis, and immune signaling. I. Pathway analysis of upregulated biological processes in stromal subpopulations, stratified by proliferative and secretory. Showing Normalized Enrichment Score (NES), p-adj <0.05 with selected pathways highlighted in bold.

The top upregulated DEGs included genes related to heat shock proteins (HSPs), specifically isoforms of *HSP70* - known to protect cellular damage caused by stress and inflammation,^19,20^ as well as for their functions as steroid hormone receptor chaperones, cell cycle regulators, and regulators of apoptosis^21^. These include *HSPA6* (adj. p-val <0.001, log2FC = 0.77 vs 2.88 for proliferative and secretory phases, respectively) and *CRYAB* (adj. p-val <0.001, log2FC = 1.36 for proliferative; adj. p-val < 0.001, log2FC = 2.48 for secretory phase). The top downregulated genes included *SFRP4* (adj. p-val< 0.001, log2FC = -1.04 for proliferative and adj. p-val<0.001, log2FC = -1.54 for secretory phase) and *SFRP1* (adj. p-val<0.001, log2FC = -0.44 for proliferative, adj. p-val<0.001, log2FC = -1.04 for secretory phase) that encode these frizzle related proteins (SFRP4, SFRP1), inhibitors of Wnt signaling^22^. The remaining 48 overlapping DEGs exhibited phase-dependent expression patterns with opposing alteration directions between the proliferative and secretory phases (**SFig. 9A**).

We applied Gene Set Enrichment Analysis (GSEA) to understand the pathway changes implicated in endometriosis versus pristine controls in the proliferative and secretory phases independently (**Fig. 3C, STable 14**). Our results showed that in the secretory phase, pathways related to heat response and protein folding were upregulated, which could represent reactive oxygen species (ROS) accumulation in this phase in disease versus control. Other apoptotic and programmed cell death pathways were additionally identified in endometriosis, as well as enrichment related to response to metal ion (**STable 14**).

### Endometrial stromal fibroblast subpopulations associated with endometriosis

To further explore these phase-specific disease phenotypes in more detail, we sub-clustered stromal fibroblasts and VSM populations and annotated them based on canonical markers, which resulted in eight subpopulations (**Fig. 3D-E, STable 2, Methods**). Among the identified fibroblast clusters, we found activated fibroblast cells, a proliferating cluster, and a previously described SFRP4+ population in endometrial fibroblasts^23^. Additionally, other distinct stromal fibroblast clusters were identified, with IGF1+, MMP+, HSP+, and WFDC2+ markers. Each subpopulation had different distributions related to the menstrual phase and disease status (**SFig. 9B-D**). All subpopulations were observed across all libraries (**SFig. 9E-F**).

We compared cell type composition between endometriosis and pristine controls, stratifying samples between secretory and proliferative menstrual phases (**SFig. 9G**). During the proliferative phase, the SFRP4+ subpopulation was less abundant in endometriosis patients (p-adj. < 0.009) (**Fig. 3F**). This cell population has been described to be abundant during the endometrium’s regenerative phase^23,24^.In the secretory phase, the fraction of HSP+ stromal subpopulation was significantly higher in cases versus controls (p-adj. < 0.0001) (**Fig. 3F**). This HSP+ population has not been previously observed in endometrial fibroblast tissue. However, it has been described in other fibroblast tissues under stress conditions^25^. Their role may be significant in collagen production and fibrosis processes^26,27^.

Furthermore, the differences between secretory and proliferative proportions were significant in activated fibroblasts (p-adj. < 0.05). Activated fibroblasts drive wound healing and are linked to cancer through their contractile, ECM-producing, and inflammatory roles^28^. The same difference in proportion was observed in the HSP+ population (p-adj. < 0.01) in cases (**SFig. 9H**).

At the transcriptional level, phase-specific differential expression was observed in several subpopulations when comparing cases to pristine controls (**Fig. 3G, STable 8,9, SFig. 10-11)**. We investigated the activated fibroblast population, which during the proliferative phase exhibited the most differences compared to pristine controls (adj. p-val < 0.05 and |log2FC|> 0.25, n=100 DEG) (**Fig. 3G**). The top upregulated genes include *TANGL* (adj. p-val < 0.01, log2FC = 1.87) and *CAV1* (adj. p-val < 0.001, log2FC = 1.26), both biomarkers of cancer-associated fibroblasts^28–30^. *NFKBIA* (adj. p-val < 0.001, log2FC = 1.25), a regulator of the inflammatory response, and *KLF6* (adj. p-val < 0.001, log2FC = 1.05), which regulates proliferation and apoptosis of endometrial stromal cells^31^, were also significantly upregulated.

Conversely, the downregulated genes included *IGFBP5* (adj. p-val < 0.001, log2FC = - 1.92), implicated in early responses during the development of fibrosis^32^, and *PENK* (adj. p-val < 0.05, log2FC = -1.74), associated with the induction of apoptosis^33^ (**Fig. 3H**).

The HSP+ population, also included differences in transcriptomic signature endometriosis compared to controls, specifically during the proliferative phase (adj. p-val < 0.05 and |log2FC| > 0.25, n=48 DEG) (**Fig. 3G,H**). Downregulated genes included *IGFBP5* (adj. p-val <0.001, log2FC=-1.66) and *SFRP4* (adj. p-val <0.001, log2FC=-1.66). Conversely, upregulated genes included *GADD45B* (adj. p-val <0.01, log2FC=1.05), which relates to stress response and immune regulation^34^, *TCIM* (adj. p-val <0.001, log2FC=0.67), a positive regulator of the Wnt/beta-catenin signaling pathway^35^, as well as *KLF6* (adj. p-val <0.001, log2FC=0.56) and *NFKBIA* (adj. p-val <0.001, log2FC=0.68), which were also observed in activated fibroblasts. Other important upregulated genes that play a role in immune response^36^ were *FOS* (adj. p-val <0.001, log2FC=0.63) and *JUN/JUNB* (adj. p-val <0.001, log2FC=0.56 and log2FC=0.56, respectively) (**Fig. 3H**). GSEA revealed that this HSP+ subpopulation exhibited an increase in immune response development and cellular response to stress (adj. p-val < 0.05) (**Fig. 3I**).

Proliferating fibroblasts also exhibited notable differences during the proliferative phase in cases compared to controls. This subpopulation showed upregulation of genes including *CCN2* (adj. p-val<0.001, log2FC= 0.76) (**SFig. 11F**), which plays a critical role in cell proliferation, differentiation, and adhesion.

We next aimed to investigate the VSM cells. This population clustered closely with the stromal fibroblast population, perhaps due to their shared mesenchymal lineage and other biological similarities. For instance, VSM cells can convert into myofibroblasts during wound healing and fibrosis, and vice versa, stromal fibroblasts are also able to transform into myofibroblasts in inflammatory environments^37^. In addition to VSM, we also examined WFDC2+ fibroblasts that express WFDC2, a marker in cancer-associated fibroblasts^38^; additionally, this gene has been shown to promote fibroblast-myofibroblast transition in pulmonary cells^39,40^. Given this similarity, we hypothesize both cell types may play shared roles in the development of endometriotic lesions.

In the secretory phase, VSM exhibited the highest number of DEGs in cases compared to pristine controls (adj. p-val < 0.05 and |log2FC| > 0.25, n=122 DEG). (**Fig. 3G**). WFDC2+ fibroblasts (WFDC2+ F) also exhibited a significant number of DEGs (n=49 DEG) (**Fig. 3G**).

Both subpopulations had upregulated DEGs involving heat shock-related proteins that protect cells from stress-induced damage. These included *HSPA6* (adj. p-val <0.001, log2FC = 2.23 in VSM) (adj. p-val <0.001, log2FC = 3.85 in WFDC2+ F), *HSPA1B*, (adj. p-val <0.001, log2FC = 1.93 in VSM) (adj. p-val <0.001 log2FC =2.99 in WFDC2+ F) and *CRYAB* (adj. p-val <0.001, log2FC = 1.43 in VSM) (adj. p-val <0.001, log2FC = 2.46 in WFDC2+ F.) (**Fig. 3H**). Downregulated genes in both subpopulations included *SFRP1* (adj. p-val <0.001, log2FC = - 0.80 in WFDC2+ F), (adj. p-val <0.001, log2FC = -0.41 in VSM.), whose suppression may promote cell proliferation and migration^41^ (**Fig. 3H**). Consistently, GSEA showed the upregulation of heat-stimulated pathways in both subpopulations and a common response to stress. Additionally, as a result of this heat shock protein-related signature, VSM showed enrichment of distinct patterns of regulation of apoptosis, including negative regulation of cell death and programmed cell death, as well as immune response, showing regulation of cytokine production (**Fig. 3I, STable 14**).

### Increased motility and migration in unciliated epithelial cells during the secretory phase

Since unciliated epithelial cells consistently exhibited high transcriptional changes in cases, we compared their expression patterns stratified by proliferative and secretory phases (**Fig. 4A, B, STable 3**). Genes that were consistently upregulated throughout the menstrual cycle in cases included those related to immunomodulation, e.g. *PAEP* (*progesterone-associated endometrial protein*) (adj. p-val <0.001, log2FC = 2.59 and adj. p-val <0.001, log2FC = 2.29 in proliferative and secretory phases, respectively) and i*nterferon stimulated gene 15* (*ISG15*) (adj. p-val <0.001, log2FC = 0.31 and adj. p-val <0.001, log2FC = 0.78 in proliferative and secretory phases, respectively), (**Fig. 4B, SFig. 12A**). In contrast, *SOX4*, a potential negative modulator for cell proliferation^42^, was downregulated in cases in both phases (adj. p-val <0.001, log2FC = -0.32 in proliferative, adj. p-val <0.001, log2FC =-1.4 in secretory phase) (**Fig. 4B**). Significant upregulation of pathways related to cell motility and migration were noted in the secretory phase (**Fig 4C**), as well as an increased response to metal ion, similar to fibroblast cells (**STable 14**). This prompted further investigation into the specific epithelial subpopulations driving these changes.

**Figure 4:**
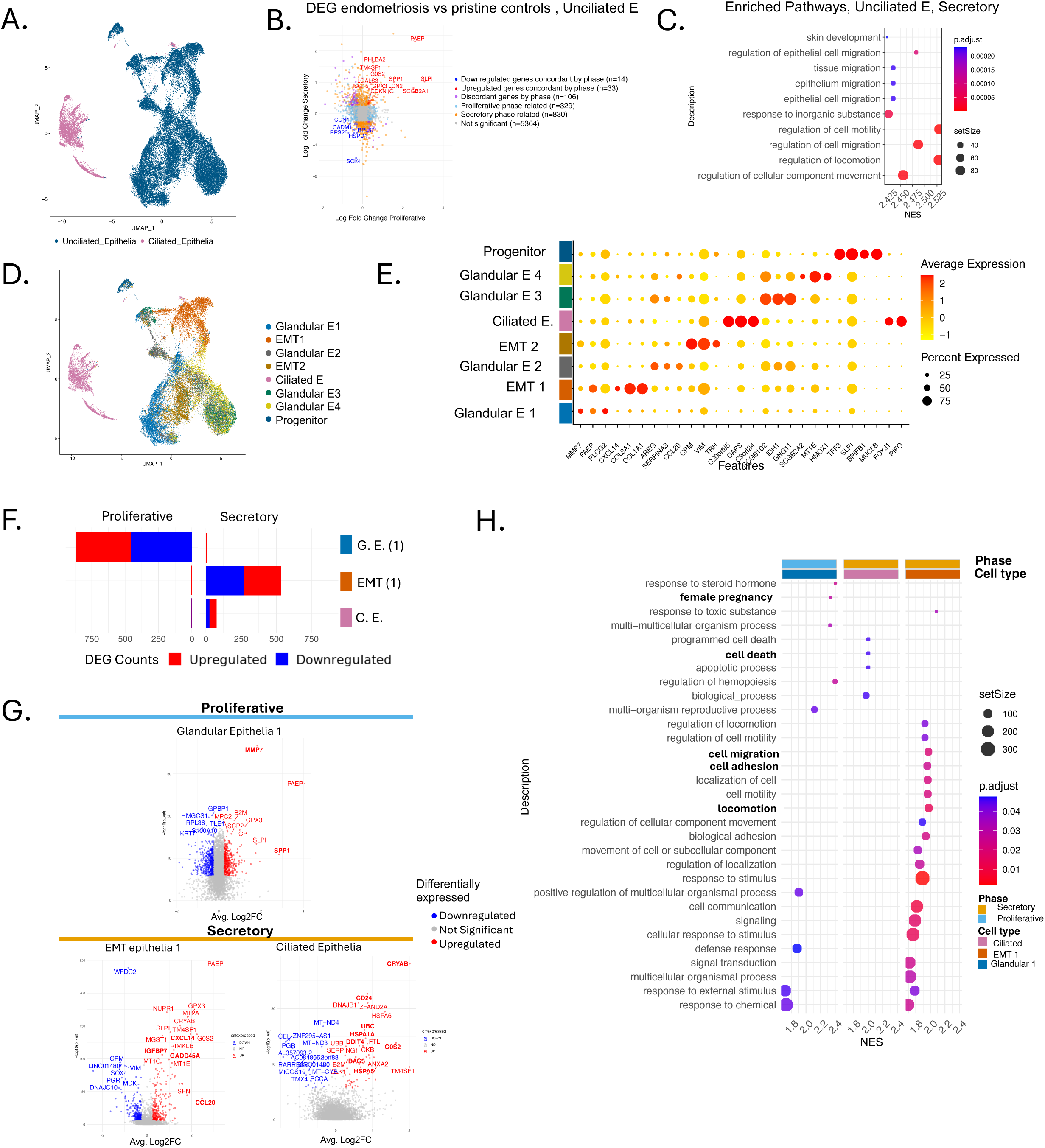
Epithelial subpopulations show upregulation of epithelium migration in endometriosis patients. A. UMAP visualization of epithelial cell subtypes, color-coded by major tissue populations (as shown in Fig. 1A). B. Log2FoldChange of DEG in the secretory and proliferative phase unciliated epithelia, comparing pristine controls and endometriosis samples. Samples were stratified by menstrual phase. DGE was assessed using a threshold of p-adj.<0.05 and |log2FC| > 0.25. C. Upregulated pathways in all unciliated epithelial populations in the secretory phase. Showing Normalized Enrichment Score (NES), p-adj <0.05. D. UMAP visualization of epithelial cell subtypes, color-coded by specific tissue type classes. E. Top marker genes for epithelial subtype. F. Number of DEGs comparing control and endometriosis samples per epithelial population stratified by menstrual phase. Red, upregulated in endometriosis, blue downregulated in endometriosis G. Differentially regulated genes between pristine controls and endometriosis samples, stratified by menstrual phase. Bolded genes are involved in selected pathways of cell death, locomotion, migration and adhesion. H. Pathway analysis of upregulated biological processes in epithelial subpopulations. Showing Normalized Enrichment Score (NES), p-adj <0.05 with selected pathways highlighted in bold.

### Endometrial glandular epithelial cells expressed genes associated with endometriosis and pregnancy

Eight distinct epithelial cell subpopulations were identified using canonical and reported markers (**Fig. 4D,E, STable 2, Methods**). These comprised four glandular epithelial populations, two epithelial-to-mesenchymal transition cells (EMT) subpopulations (one *VIM*+ and one with collagen-related gene expression), a progenitor cell population (*MUC5B*+ and *TFF3*+)^14^, and a large ciliated epithelial cell (CE) subpopulation previously mentioned.

We observed pronounced variations in subpopulation composition across menstrual cycle phases (**SFig. 12B**) and disease condition (**SFig. 12C,D**). Some subpopulations, here labeled glandular E3 and glandular E4, were composed of specific libraries or samples (**SFig. 12E-F**). Exploring differences in cell type proportion per sample showed no significant changes between endometriosis and controls (**SFig. 12G,H**).

In the proliferative phase, several epithelial subpopulations exhibited notable transcriptional differences between cases and controls (**Fig. 4F, STable 10-11, SFig. 13 ,14**). In glandular E1, we identified 870 DEGs between endometriosis and pristine controls during the proliferative phase. Genes upregulated in this subgroup were also upregulated in the larger category of unciliated epithelial cells, including expression of *PAEP* (adj. p-val <0.001, log2FC=4.06) (**Fig. 4G**). GSEA using glandular epithelium DEGs revealed distinct pathway alterations associated with endometriosis. In particular, pathways associated with “Female pregnancy” and “Response to steroid hormones” were enriched (**Fig. 4H**), and several genes belonging to these pathways were upregulated in endometriosis cases, including *SPP1, MMP7, IGFBP7, FOSB, LIF,* and *SLC2A1* (**STable 10,14**). While these genes are crucial for pregnancy and fertility, they are also implicated in the pathophysiology of endometriosis^43–48^. Additionally, a gene related to immune response in the endometrium^49^, *CD74*, was upregulated (adj. p-val <0.001, log2FC=1.09) (**STable 10**). These findings highlight the complex endometrial microenvironment in the setting of endometriosis

### EMT epithelial subtypes dominate the changes in unciliated epithelial cells

During the secretory phase, two main subpopulations exhibited differences at the transcriptional level between endometriosis patients and pristine controls: EMT1 and CE cells (adj. p-val < 0.05 and |log2FC| >0.25, n=533 DEG for EMT1 and n=75 for CE) (**Fig. 4F**). In particular, *CRYAB* (adj. p-val <0.001, log2FC=1.82 for EMT1, log2FC=2.01 for CE) was differentially expressed in both subpopulations (**Fig. 4G**). This upregulation seen in EMT1 and CE cells is also common to other cell types, such as fibroblasts and VSM. Uniquely, EMT1 showed upregulation of pathways related to cell migration, locomotion, motility, and adhesion (**Fig. 4H**). The genes driving this enrichment were *CCL20*, (adj. p-val <0.001, log2FC=2.20), *CXCL14* (adj. p-val <0.001, log2FC=1.95), *IGFBP7* (adj. p-val <0.001, log2FC=0.68), and *GADD45A* (adj. p-val <0.001, log2FC=0.98). Furthermore, we observed a decrease in the regulatory processes of the canonical Wnt signaling pathway (**STable 14**) with inhibitors *SFRP1* and *SFRP4* downregulated, suggesting activation of the WNT beta-catenin pathway. In contrast, CE showed an upregulation of pathways related to apoptosis and cell death (**Fig. 4H**). These data underscore unique phase-specific differences in epithelial subtype gene expression in endometrium from endometriosis patients.

### Endothelial cells show leukocyte migration and inflammation associated with endometriosis

Endothelial cells exhibited a significant number of DEGs in cases versus controls across menstrual cycle phases (**STable 3**). Notably, genes involved in vascular function and inflammation, such as *KLF6* (adj. p-val <0.001, log2FC=1.77 in secretory phase; adj. p-val <0.01, log2FC=0.61 in proliferative phase), *ICAM1* (adj. p-val <0.001, log2FC=1.79 in secretory phase), and *APOLD1* (adj. p-val <0.001, log2FC=0.62 in secretory phase), were upregulated in endometriosis cases (**SFig. 6G,H**). GSEA again revealed upregulated pathways related to heat shock protein response and oxidative stress in the secretory phase, emphasizing the impact of endometriosis on these key biological processes across cell types. Moreover, enrichment of leukocyte migration was noteworthy in the secretory phase, alongside regulation of cytokine expression in both menstrual phases **(SFig. 7F, STable 14**), thus highlighting the interaction between endothelial cells and immune regulation.

### Immune response in macrophages and NK cells in endometriosis

Prior work has described the critical role of the immune system in maintaining endometrial homeostasis, with immune dysregulation contributing to endometrial pathology and pregnancy complications^50,51^. Given the heterogeneity of immune cells, we annotated the broad immune population into 11 distinct subsets using canonical immune cell markers: macrophages, monocytes, conventional type 1 dendritic cells (cDC1), plasmacytoid dendritic cells (pDCs), mast cells, uterine natural killer (NK) cells, innate lymphoid cells, plasma cells, stressed T cells^52^, CD8+ T cells, T regulatory (Treg) cells. Additionally, a subset of lymphatic endothelial cells was observed to cluster with the immune cell population (**Fig. 5A,B**, **STable 2)**. The distribution of these cell types varied across disease conditions and menstrual phases (**SFig. 15A-C**). Additionally, the cell composition varied across patients and libraries **(SFig. 15D-E).** An exploration of differential abundance revealed no significant differences between disease and pristine control or among menstrual phases in immune cell populations (**SFig. 15F-G**).

**Figure 5:**
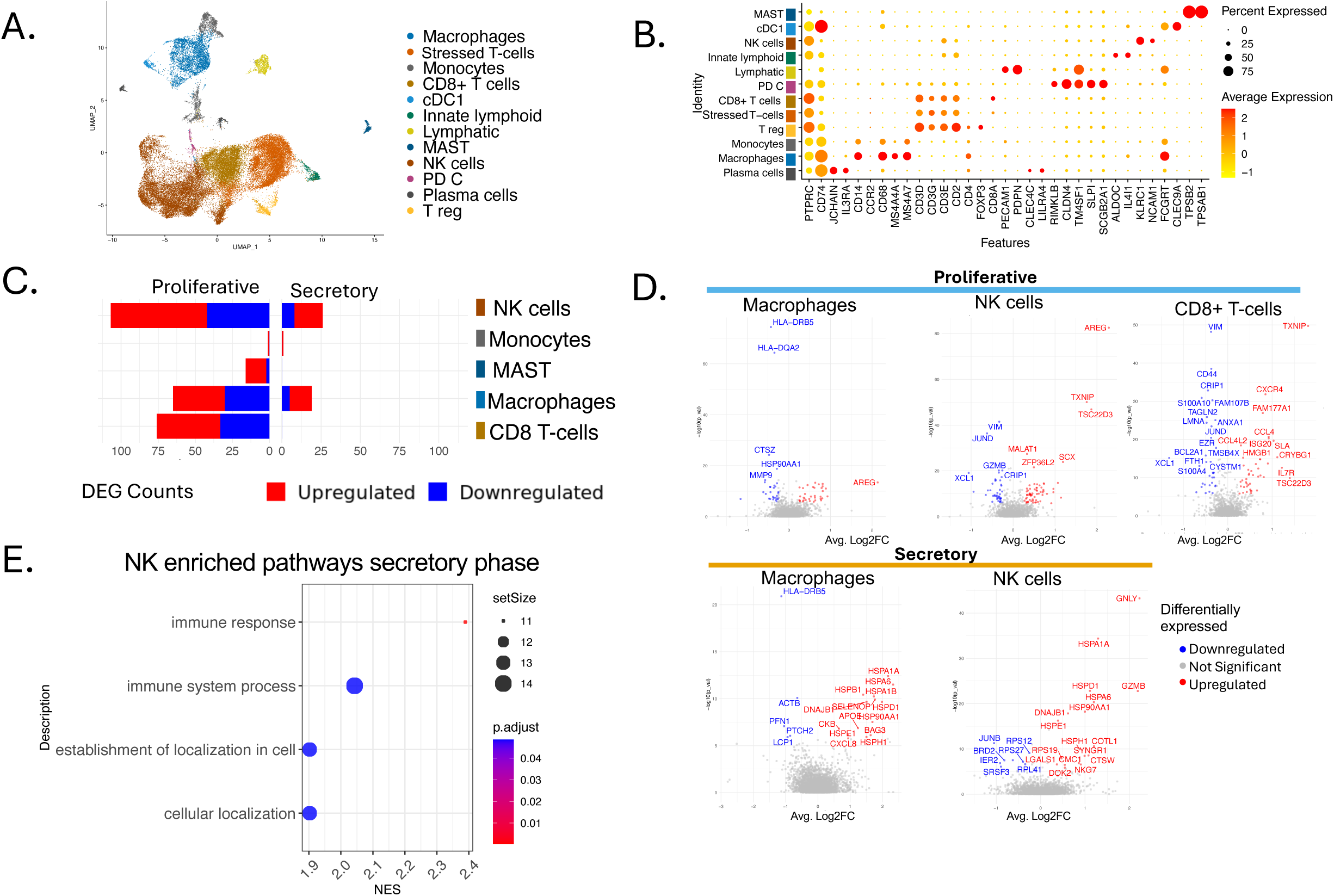
Uterine NK cells demonstrate augmented activity during endometriosis. A. UMAP overview depicting fine-grained immune subtypes. B. Selected marker genes for each subpopulation. C. Number of DEGs comparing pristine control and endometriosis samples per immune population, stratified by menstrual phase. Red, upregulated in endometriosis, blue downregulated in endometriosis. D. Differentially regulated genes in macrophages, NK cells, and CD8+ T-cells, between pristine controls and endometriosis samples, stratified by menstrual phase. E. Pathway analysis of upregulated biological processes in NK cells. Showing Normalized Enrichment Score (NES), p-adj <0.05.

Differential gene expression analysis of immune cell subsets identified macrophages, NK cells and CD8+ T cells as the populations with the most transcriptomic differences between endometriosis versus controls (**Fig. 5C, STable 12,13**). In the secretory phase, macrophages in endometriosis eutopic endometrium samples exhibited upregulation of HSP genes related to stress-induced damage protection (*HSPA1A* adj. p-val <0.001, log2FC = 2.17, *HSPA6* adj. p-val < 0.001, log2FC = 2.32) and cytokine-related genes (*CXCL8*, adj. p-val < 0.05, log2FC = 0.93). Additionally, macrophages in endometriosis showed downregulation of Class II HLA genes, which encode human leukocyte antigen critical for immune recognition, in both the proliferative (*HLA-DQA2*, adj. p-val < 0.001, log2FC = -0.34, *HLA-DRB5*, adj. p-val < 0.001, log2FC = -0.43) and secretory phases (*HLA-DRB5*, adj. p-val *<* 0.001, log2FC=-1.13) (**Fig. 5D**). This downregulation of HLA genes also occurred in conventional dendritic cells (cDC) during the proliferative phase (*HLA-DRB5*, adj. p-val < 0.001, log2FC=-0.88; HLA-*DQA2* adj. p-val < 0.001, log2FC=-1.2) (**STable 12**).

CD8+ T cells also exhibited transcriptional differences in endometriosis patients, including upregulation of *TXNIP* (adj. p-val < 0.0001, log2FC=1.81) (**Fig. 5D**), which acts as a regulator of cellular oxidation^53^. In addition, *XCL1* (adj. p-val < 0.0001, log2FC=-1.32), which is expressed as part of the inflammatory response and *ISG15* (adj. P-val < 0.0001, log2FC=-0.71), recognized for its immunomodulatory properties, were both downregulated in CD8+ T cells in endometriosis, suggesting impaired immune surveillance.

NK cells exhibited many differences in both proliferative and secretory phases. In this population, genes related to cytotoxic activity and chemokine-related recruitment of other immune subsets (*GZMA* adj. p-val < 0.001, log2FC=-0.48, *GZMB* adj. p-val < 0.001, log2FC=-0.34, *XCL1*, adj. p-val < 0.0001, log2FC=-1.06) were downregulated in cases during the proliferative phase, while *TXNIP* was upregulated (adj. p-val < 0.001, log2FC=1.75) (**Fig. 5D, STable12**). However no specific pathways were upregulated in the proliferative phase.

In the secretory phase, we detected overexpression of HSPs (*HSPA1A* adj. p-val < 0.001, log2FC=1.30, *HSPA6* adj. p-val < 0.001, log2FC=1.20). Notably, *GNLY*, which promotes the induction of the immune response, was upregulated in this phase (adj. p-val < 0.01, log2FC = 2.22), along with the cytotoxic gene *GZMB* (adj. p-val < 0.001, log2FC=2.19) (**Fig. 5D, Stable 12**). This pattern was also observed in the enrichment of pathways using GSEA, which showed increased immune response and immune system processes in NK cells during the secretory phase (**Fig. 5E, Stable 14**).Together, these findings suggest a specific impact of endometriosis on uterine NK cell physiology during the secretory phase, which may not only affect fertility, but also risks of pregnancy complications and tissue regeneration in the absence of pregnancy, given the proposed roles of this specialized cell type.

### Inferred communication between immune and non-immune cells in endometriosis

To gain insight into the interplay between immune and non-immune cells, beyond transcriptional changes. To explore this, we used CellChat^54^ to compare cases versus pristine controls, stratified by phase. Overall, we found distinct inferred receptor-ligand communication patterns in both phases (**Fig. 6A**). In the proliferative phase, we identified a mix of upregulated and downregulated pathways, including interactions exclusive to either cases or controls. In contrast, in the secretory phase, we observed mainly upregulation of pathways in endometrium from cases (**Fig. 6A**).

**Figure 6:**
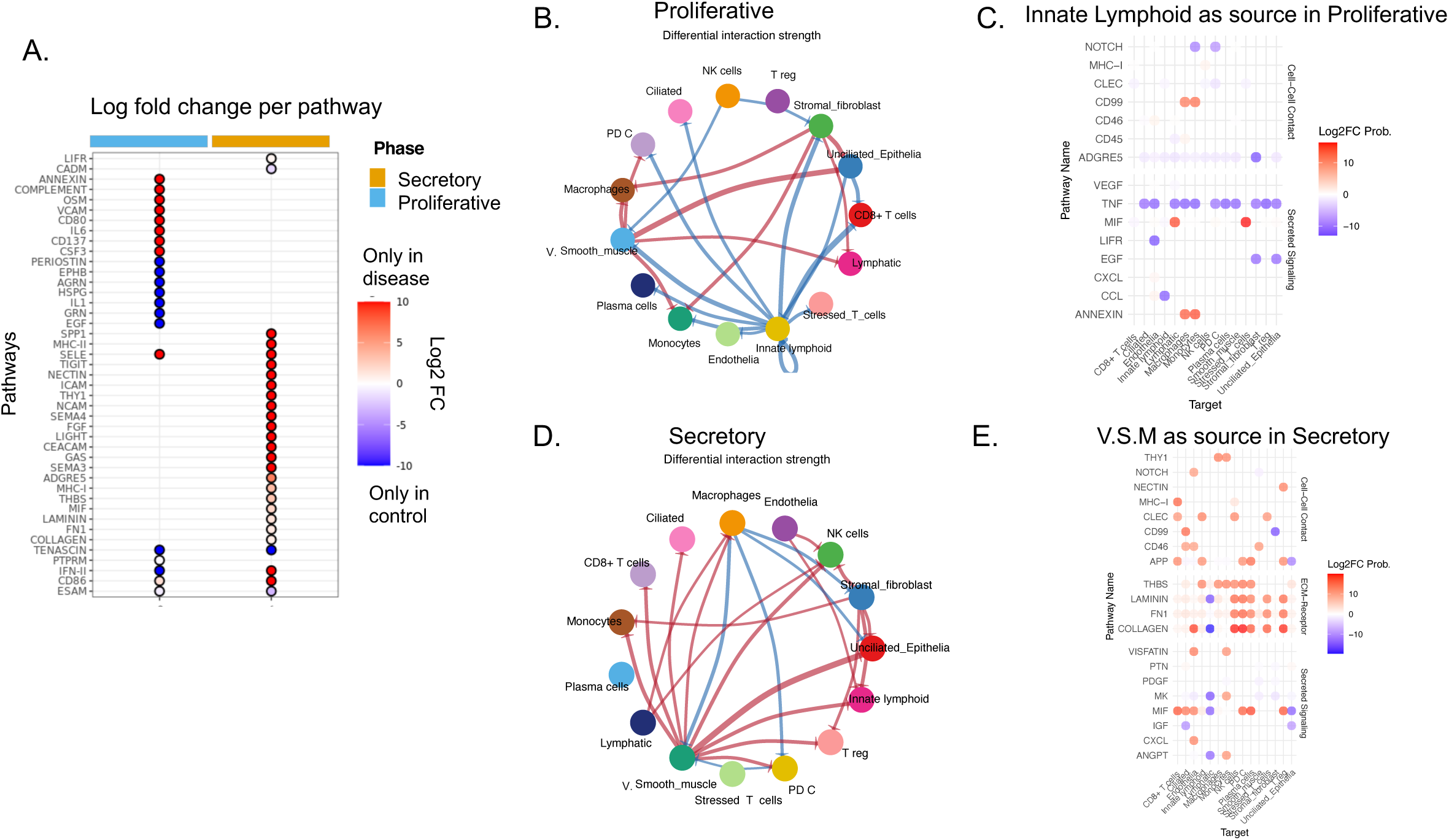
In-silico prediction shows upregulated communication in extracellular matrix and inflammatory pathways in disease. A. Summary of differential predicted pathways stratified by phase, color by log2-fold change and stratified by menstrual phase. B. Pathway differences in the proliferative phase. P-value < 0.05, top 10% of interactions. Red represents increase in endometriosis compared to pristine controls; blue represents decrease. C. Probability of communication with Innate lymphoid cells as the source in the proliferative phase, colored by Log2FC differences between endometriosis and controls. D. Pathway differences in the secretory phase. P-value <05, top 10% of interactions. Red represents increase in endometriosis compared to pristine controls; blue represents decrease. E. Probability of communication with VSM cells as the source in the secretory phase, colored by Log2FC differences between endometriosis and pristine controls.

To investigate the most significant differences in communication pathways by cell type, we visualized the top 10% differentially inferred pathways, stratified by phase. In the proliferative phase, we observed that innate lymphoid cells (ILCs) exhibited the most significant communication differences, characterized by decreased signaling to both immune and nonimmune populations in cases (**Fig. 6B**). By identifying specific communication pathways, we observed that ILCs in cases sent a decreased signal in TNF pathways (**Fig. 6C**). Further analysis of ligand-receptor pairs in this population revealed that *TNFRSF1A* and *TNFRSF1B* were only present in control samples (**SFig. 16**). While, TNF ligands are recognized for their role in regulating inflammatory pathways in in endometriosis^55^, it is hypothesized that increased levels of soluble TNF receptors may serve as a regulatory mechanism to sequester and neutralize TNF in various chronic inflammatory conditions^56^. Notably, TNFRSF1A has been identified as a prospective therapeutic target for endometriotic lesions based on in vivo studies. studies^57^.

We performed a similar analysis in the secretory phase (**Fig. 6D**). VSM and stromal fibroblasts exhibited the most upregulated interactions, acting as key ligand sources targeting immune cells, including NK cells, CD8+ T cells, and macrophages. To investigate which pathways where VSM were the sources of upregulation, we compared the differences in signaling probability between cases and control samples (**Fig. 6E**). These highlighted the *THBS* glycoprotein pathway and *FN1* pathway, both of which involve extracellular matrix (ECM) proteins and interact with cell membrane receptors. Another important pathway was *COLLAGEN*, with a presence in both controls and cases but significantly upregulated in endometriosis. Collagen is an important ECM-interacting protein. One of its potential roles in endometriosis is its deposition in endometriotic lesions as a result of chronic inflammation^58^. However, its mechanistic function in the endometrium of patients with the disease remains to be determined.

## Discussion

Endometriosis is a prevalent and complex disease and understanding its impact on different cell subtypes of the eutopic endometrium as its tissue of origin and as a target of systemic associated with this disorder remains a challenge. It has been reported that eutopic endometrium differs between individuals with and without endometriosis^59–61^ .To investigate this, we leveraged a comprehensive data set, containing over 235,000 cells encompassing people with endometriosis, fibroids, and without disease across the menstrual cycle.

Studies have demonstrated that hormonal responses, especially to sex steroid hormones during the menstrual cycle, affect endometrium and also endometriosis lesions^1,62,63^. This emphasizes the need to obtain and analyze samples by menstrual phases, and focus on samples with no additional uterine/endometrial pathology and without exogenous hormonal exposures^20^. By including samples spanning the menstrual cycle, avoiding hormonal interventions, and incorporating pristine controls, we were able to minimize potential sources of bias and better isolate disease-specific signals.

As we have shown previously^64^, we established that the menstrual cycle phase influences the epigenomic and transcriptomic endometrial landscapes. Herein, we have found that specific cell types, such as fibroblasts and epithelial cells, contribute to endometrial pathophysiology associated with endometriosis through mechanisms involving oxidative stress and aberrant apoptosis signaling. We also demonstrated that alteration of immune cell communication contributes to these processes. This study provides a valuable map of biological pathways that characterize endometriosis in the eutopic endometrium across menstrual cycle phases.

Building on this rationale, our study suggests that the differences between endometriosis and controls are heavily dependent on cycle phase. Ours and other studies have demonstrated that endometrial samples cluster more strongly by menstrual cycle phase than disease status at molecular and epigenetic levels^61,65,66^. Consistent with these findings, our PCA revealed distinct clustering related to the menstrual cycle phase independent of disease status. Furthermore, we also showed consistent phase-specific differences at the pathway and transcriptomic levels between endometriosis cases and fibroid controls, and pristine controls.

This phase-specific response is further supported by our observations of an increased expression of heat shock protein (HSPs), especially HSP70-related isoforms^67^, in secretory-phase endometrial samples. HSP70 is considered a marker of oxidative stress in endometriosis in lesions and in the eutopic endometrium^19,68^. HSPs influence inflammation pathways, resulting in intrinsic apoptosis^69^. In fact, single-cell technology has identified HSPA6-related fibroblasts with a pro-inflammatory phenotype in various tissues^25^.

Several studies indicate elevated levels of HSPs in both eutopic and ectopic endometrial tissues compared to controls^68,70–72^, although the relative increase between these tissues requires further research. The phase-specificity of this increase also remains contentious, with some research suggesting phase-independent elevation, while others report a rise specifically during the secretory phase^71,72^. Furthermore, differing mechanisms may underlie this increase, depending on whether the tissue is eutopic or ectopic, and perhaps patient-level factors that are unique to their history of environmental/medication exposures^70^. This pattern of high HSP expression could be explained by the elevated reactive oxygen species, lower antioxidant levels, and disrupted iron metabolism that occurs the in endometriosis disease process. Endometrium from cases revealed that stromal fibroblasts and unciliated epithelia in the secretory phase had elevated metal ion-related processes, accompanied by reduced apoptosis signals, suggesting that endometrial cells may regulate ferroptosis susceptibility through oxidative pathways to support growth and proliferation, as previously observed in ectopic lesions ^70,73,74^.

Elevated oxidative expression was also present in specific cell types, including WFDC2+ fibroblasts, uterine NK cells, macrophages and ciliated cells. Previous studies have demonstrated that macrophages secrete HSP70-inducing proinflammatory cytokines *in vitr*o in the context of other diseases^75^, which was also observed in our results. Furthermore, this oxidative pattern was consistent with *TXNIP* expression in uterine NK and CD8⁺ T cells in endometriosis samples. This expression is also required for the cleavage of HSP90^76^, potentially contributing to the pro-inflammatory microenvironment of endometrium in patients with endometriosis^53,77^,. Another HSP-related response was the upregulation of CRYAB, which was consistently upregulated in endometriosis across all broad cell populations in the secretory phase. This gene has been implicated in cellular stress and also in decidualization^78,79^.

Interestingly, therapies targeting oxidative stress have been proposed for the treatment of endometriosis. Notably, TNFR1 signaling has emerged as a potential therapeutic approach^70,80^. This is particularly compelling given our findings of TNFR1 downregulation in endometriosis cases, as revealed by our cell–cell communication analysis.

The accumulation of HSPs in endometriosis also suggests cellular proliferation, promoting the survival and implantation of cells at ectopic sites^81^. This pathologic implantation is aided by reduced apoptosis in endometrial cells, allowing these cells to evade clearance mechanisms and continue growing in ectopic lesions^82,83^. This is also confirmed by our finding of increased negative regulation of cell death among fibroblasts, VSM and CE.

Besides enhanced cellular proliferation and reduced cell death, increased cell migration is another potential mechanism for disease establishment originating in the eutopic endometrium and continued growth of endometrial-like tissue ectopically. Notably, the aberrant activation of Wnt/β-catenin pathway in the menstrual endometrium has been documented in endometriosis patients and it may worsen the disease by increasing cell migration and invasion. In our results we observed a consistent downregulation of *SFRP4* and *SFRP1*. These genes inhibit Wnt signaling by preventing the interaction with WNT ligands^84^. The downregulation of these inhibitors supports the abnormal activation of this pathway^22^. We also showed that WNT regulation is decreased at the pathway level in EMT1 cells.

In our analysis, we identified two distinct EMT populations. One cell population exhibited mesenchymal and epithelial markers (EMT1). By contrast, the second EMT population showed expression of *VIM*+ as a marker (EMT2). Notably, during the secretory phase, EMT1 population showed increased cell migration, locomotion, cell motility, and cell adhesion in endometriosis. These findings align with previous research indicating that EMT is an important pathogenic mechanism, by losing its polarized cytoskeletal organization and cell-to-cell connections and increasing motility^85,86^. EMTs are stimulated by estrogen levels and hypoxia signals in eutopic and ectopic endometrium^86,87^, consistent with our finding of high oxidative stress. Additionally, a key regulation of EMT mobility, vimentin, was downregulated in this population. This encourages further exploration, as previous studies have reported conflicting results regarding the significance of VIM and other key EMT markers^86,88–90^, showing differences between eutopic and ectopic endometrium for master EMT genes^88^.

Cell migration is significantly influenced by the dynamic interactions between cells and their extracellular matrix (ECM)^91^. Aberrant expression of ECM-related factors was shown in our cell-cell communication analysis, with ECM interactions upregulated in VSM, including ligands for collagen and FN1. Furthermore, in the proliferating fibroblast population, we observed an upregulation in CCN2, which coordinates key components of the ECM, tissue repair, and growth factors^92^. This highlights the significance of ECM-related genes in endometriosis^45,93,94^. This was also shown by the upregulation of *MMP7* in glandular epithelia. MMPs in endometriosis could influence processes such as invasion, angiogenesis, inflammation and fibrosis, and are exacerbated by oxidative stress^89,95^.

The role of immune cells in endometriosis is pivotal for the disease’s progression, for example, through their interactions with non-immune tissue types. This was also observed in endothelial cells, where leukocyte migration was upregulated, along with genes related to this process for example *ICAM1*^96^. This process is essential for the induction, maintenance, and regulation of immune responses and can be influenced by cytokine expression, which affects the functionality of other cell types in preparing for and sustaining pregnancy, as well as maintaining overall tissue homeostasis.

Another critical component in the progression and clinical management of endometriosis is immune surveillance and regulating inflammatory response. Immune recognition is influenced by MHC class II molecules, expressed by antigen-presenting cells like macrophages, which are encoded by HLA genes that bind various peptides^97^. Our results show a reduction in *HLA-DR* expression in the macrophage and cDC populations in cases, indicating a functional impairment in antigen-presenting cells. This reduction was also found in previous studies of women with infertility, both with and without endometriosis^98^. Further research is needed regarding the role of the HLA system in this disease^99^. We also observed downregulation of chemokine *XCL1* in CD8+ T cells and NK cells in the proliferative phase, which play a critical role in recruiting other immune cell types such as DCs^100^.

Interestingly, we found contradictory expressions of specific cytotoxic molecules in NK cells, including upregulation of *GNLY* in the secretory phase and phase-specific expression of *GZMB* in samples from cases versus controls. The change in expression of these genes in endometriosis has been debated, with evidence for both downregulation and upregulation^101,102^. The observed upregulation of *GNLY* could be due to the inflammatory conditions of the endometrium. Conversely, the downregulation of *GZMB* and *GZMA* could suggest blunted NK cell activity, which may facilitate the progression of endometriosis. In general, this result highlights how endometriosis involves a complex interplay of inhibitors and activators.

An unexpected finding was the upregulation of *PAEP* in endometriosis samples, also known as glycodelin (Gd) in endometrial epithelial cells. Gd is thought to be involved in establishing endometrial receptivity and is considered a biomarker of the window of implantation^103,104^. Single-cell analysis further associated *PAEP* expression with a distinct epithelial sub-cluster found exclusively in control samples, opening the window of implatantion^15^. However, studies have also shown mixed results regarding Gd levels, with some reporting elevated serum levels linked to disease infiltration, while others found only minor differences in expression between control and endometriosis^105–108^. In our analysis, this gene was persistently upregulated in endometriosis samples in both proliferative and secretory phases in the unciliated epithelial population. We propose that this acyclic, constitutive expression may reflect dysregulation of this receptivity marker and immune modulator in the eutopic endometrium, making it unfavorable for embryo implantation. Discrepancies among multiple studies may be attributed to variations in sampling time points. Gd expression fluctuates throughout the menstrual cycle, peaking during implantation. However, in endometriosis, altered progesterone response could lead to a different pattern of Gd expression, potentially causing fertility issues^109^. Future research focusing on more precise time points of the menstrual cycle could help answer these questions.

In this study, we present a wide-ranging analysis of patients with endometriosis, alongside patients with uterine fibroids and pristine controls. Although our pristine controls (ovum donors not undergoing ovarian stimulation) did not undergo surgical confirmation to rule out endometriosis, it is important to note that they had no history of gynecologic disorders or symptoms of endometriosis.

While we expand upon previous studies by including samples from both the proliferative and secretory phases, the current study’s limited sample size may restrict the ability to detect subtle phase-specific changes. Including additional time points to create a more continuous map of the menstrual cycle, as well as a better representation of the late and early proliferative and secretory phases, would strengthen our findings. We found limited detection of differential abundance across cell types. We suggest increasing the sample size, especially for these time points not captured, which would provide greater statistical power. Additionally, while the inclusion of multiple endometriosis stages could help characterize the disease, the heterogeneity of sample stages and types may complicate the identification of specific differences between endometriosis variants.

Overall, our results highlight the importance of oxidative stress, apoptosis, and extracellular matrix organization as potential contributors to the pathophysiology of endometrial dysfunction in endometriosis patients. Additionally, we have identified key players such as EMT cells, specific endometrial fibroblast populations, and altered communication in NK cells and macrophages in this disease. These discoveries enhance our understanding of the complex interplay in the eutopic endometrial microenvironment in patients with endometriosis. Moreover, we provide multiple comparative analyses, examining not only the differences between endometriosis and healthy controls but also considered various stages of endometriosis and fibroid controls. We observed a notable similarity between fibroids and endometriosis, underscoring the critical importance of utilizing robust control groups.

Furthermore, understanding these eutopic alterations is crucial not only for the pathogenesis of lesions but also for helping patients experiencing fertility problems. Identifying transcriptomic signatures in eutopic endometrium could pave the way for less invasive diagnostic approaches to endometriosis. For example, our data could contribute to novel diagnostic strategies through transcriptomic prediction and drug interaction studies, and it could be leveraged for the development of targeted mono-or multi-modal therapies. We believe this study represents a valuable resource that serves as an atlas to better understand the impact of endometriosis, particularly on the eutopic endometrium.

## Methods and Materials

### Subject details

This study was conducted in accordance with the Institutional Review Board (IRB) guidelines of the University of California, San Francisco under an approved IRB protocol (#10-02786) and in accordance with the Declaration of Helsinki. Informed written consent was obtained from all subjects before endometrial tissue sampling. Subjects were 20-45 years old with regular menstrual cycles (28–30 days) and body mass index 19–29 kg m2 (inclusive) (**STable 1**). Cases included subjects with rASRM stages I-IV, endometriomas, superficial, and/or deep infiltrating disease diagnosed at surgery and confirmed histologically. Controls included samples from subjects without endometriosis but undergoing surgery for symptomatic uterine fibroids/ leiomyoma, a common condition of reproductive aged women^110^ (“pathologic controls”) and from healthy ovum donor volunteers (“pristine controls”). Subjects were excluded if they had used hormonal contraception or GnRH analogue less than 3 months prior to sampling, had polycystic ovary syndrome, and reproductive tract infections or malignancies. Some endometriosis cases additionally presented adenomyosis, fibroids, or uterine polyps (**STable1**). Cycle phase was determined by endometrial histology^111^, serum estradiol and progesterone levels, and last menstrual period. Biospecimens were obtained and processed according to the UCSF Uterine Disorders Tissue Biorepository standard operating procedures^112^.

### Endometrium tissue dissociation, cryopreservation, and epithelial cell enrichment

A two-stage dissociation protocol was used to dissociate endometrial tissue and separate it into stromal- and epithelial-enriched single-cell suspensions. Undissociated endometrial tissue was rinsed with PBS (Sigma) to remove blood and mucus. Rinsed tissue was minced into the smallest possible pieces (∼1×1×1 mm) with disposable scalpels on a petri dish and dissociated in collagenase I or IV (Sigma) at 37 °C for 1hr in a 15 ml Falcon tube with constant agitation on a Nutator. This primary enzymatic step dissociates stromal cells into single cells while leaving epithelial glands and lumen mostly undigested. At this stage, the epithelial portion was either separated from the stromal cells via a 40-μm cell strainer or left undigested within the stromal cells. If present, red blood cells were lysed with 1× RBC Lysis Buffer (Invitrogen). The resulting suspensions were then cryopreserved using DMSO-based cryoprotectant (80% DMEM, 10% serum, 10% DMSO) following standard protocol.

The supernatant was therefore collected as the stromal fibroblast-enriched suspension. The epithelial pellet was dissociated in 1 ml Accutase (Millipore) for 30 min at 37 °C, during which homogenization was performed every 15 minutes via intermittent pipetting using a 1mL pipette. DNaseI (Roche) was then added to the solution to a final concentration of 0.2mg/mL to digest extracellular genomic DNA and was quenched with 9 ml of DMEM after 1 min of incubation at room temperature. The resulting cell suspension was pipetted, filtered through a 40-μm cell strainer, and centrifuged at 300 RFC for 5 min. The pellet was resuspended as the epithelium-enriched suspension.

### Sample multiplexing and randomization

Multiplexing via natural genetic variants^113^ was used for single-cell capture, cDNA generation, and library preparation. Briefly, samples from 4 different individuals were pooled and loaded onto a single well of the Chromium Next GEM Chip G (10x Genomics). For demultiplexing, an aliquot of each sample was analyzed separately by bulk RNAseq to associate high-confidence genetic variation with individual samples of origin.

For multiplexing, stratified randomization^114^ was used to create groups of 4 individuals to balance the influence of the following covariates: conditions (pristine controls, pathological controls, stage I-II endometriosis, stage III-IV) and menstrual cycle phase (proliferative, secretory). Briefly, samples were classified into categories based on the combination of the covariates above. A random number between 1∼n (n=desired number of groups) was assigned to each sample in each of the categories, with 4 being the maximum number of samples per group, such that each sample was assigned to n groups of 4 samples.

### Single-cell sequencing

*Single-cell capture, cDNA generation, and library preparation via Chromium 10x platform and chemistry.* A total of ∼60,000 cells from four individuals (∼15,000 cells/each) were loaded onto a single well of the Chromium Next GEM Chip G (10x Genomics). GEM generation and barcoding, reverse transcription, cDNA generation, and library construction were performed following the manufacturer’s protocol (Single cell 3′ reagent kit v.3.1, 10x Genomics). Specifically, 11 PCR cycles were used for cDNA generation, whereas 11-13 cycles were used for PCR during library preparation, depending on the amount of cDNA input and the manufacturer’s guidelines.

*Bulk RNA extraction, cDNA generation, and library preparation.* For each sample, bulk RNA extraction was performed on ∼100,000 cells using AllPrep DNA/RNA Microkit and protocol Simultaneous Purification of Genomic DNA and Total RNA from Animal and Human cells (Qiagen). The resulting RNA was diluted and quantified via Nanodrop and Fragment analyzer. 1.5ng total RNA with RIN>8 was used for cDNA generation using SMARTseq v4 chemistry (Takara) and library prepped using Nextera DNA Flex Library Preparation Kit (Illumina, San Diego, CA).

*Sequencing*. Single-cell libraries were sequenced to a depth of ∼25,000 reads per cell on a Novaseq. Bulk RNAseq libraries were sequenced to a depth of ∼30M reads per sample on a HiSeq.

### Single-cell data processing and filtering

Thirteen libraries were independently processed using the Cell Ranger^115^ software pipeline (version 3.1.0). Reads were mapped to the transcriptome using the human hg38 version, and UMI counts were calculated using default parameters. The samples were demultiplexed using Freemuxlet^113^ using the number of samples per pool and a list of valid barcodes. We processed each UMI count matrix using the R package Seurat (v.4.0.3)^116^. Using Freemuxlet cells were classified according to their UMI content and doublets, and ambiguous cells were filtered, while singlets were retained. Further filtering per each pool included removing cells with high mitochondrial content (>20%), high ribosomal content (>50%), high RNA count (50,000>), low feature (<200) and low RNA count (<300). We performed normalization and scaling. We calculated variable features, principal component analysis (PCA), and ran UMAP and clustering (Leiden). To assign demultiplexed cells to the sample of origin, genetic profiles between demultiplexed cells and bulk RNA profiles were compared using bcftools (v.1.10). Before integration we performed a second filtering step, we removed clusters which top markers were mitochondrial genes. We employed Seurat to combine the data from the 13 pools into a single object. During this process, mitochondrial, ribosomal, and cell cycle scores were regressed out during scaling. To correct batch effects, we utilized Harmony^117^ to perform pool integration. Subsequently, we conducted PCA on the variable genes (n=3000), and it was used as input for an unsupervised cell clustering approach using the Louvain algorithm. Dimensionality reduction and visualization were performed using UMAP. One individual using hormonal medication was removed after clustering. Following cell filtering, the single-cell dataset consisted of 228,021 cells with 29,001 genes expressed. (**STable 1**).

### Identification of cell types

Broad cell types were classified through a combination of manual and automated annotation. Cell type-specific markers were identified using the FindMarkers function in Seurat with default parameters. Additionally, clusters exhibiting high mitochondrial gene expression among their top 10 markers were excluded from further analysis. Subsequently, the remaining cells were re-clustered. This process was repeated until the top 10 markers for every cluster were not part of the mitochondrial family.

For reference annotations, a previous dataset of the human endometrium was utilized^118^. This dataset was downloaded, checked for quality, and their labels verified. This dataset was then used as reference using SingleR^119^ for automated annotation of our broad main cell types. This annotation was cross-referenced with the top gene markers.

*Identification of fine cell types.* Epithelial, stromal, and immune subpopulations were identified using a tailored approach with Seurat’s FindNeighbors and FindClusters functions. Each cell type was analyzed at a resolution parameter adjusted according to its heterogeneity. Identification of specific markers was subsequently performed using the FindMarkers function in Seurat with default parameters. Clusters expressing mitochondrial genes (MT) among their top 10 markers, or immune-specific genes in non-immune cell types, were excluded. The cells were then re-clustered. This process was repeated multiple times for epithelial cells until no clusters with high MT expression remained. Manual annotation of the identified clusters was conducted based on markers described in the literature. The result of these subsets were consequentially used for downstream analysis, for broad and specific cell types.

*Fibroblast Fine Annotation:* After re-clustering, we obtained seven fibroblast clusters and the previously reported vascular smooth muscle (VSM) cluster. We identified Cluster 0 (C0) with marker S100A4, described as an activated fibroblast^120^; C1 with IGF1+ expression, indicating decidualization^121^; C2 with HSP markers, suggesting stress; C3 with SFRP4+ expression, a previously described cluster^23^; Cluster C4 with matrix metalloproteinase (MMP) genes related to the extracellular matrix^122^; C5 with TOP2A+ expression, a proliferative marker^123,124^; C6 expressing VSM markers; and C7 with WAP four-disulfide core domain 2 (WFC2) expression.

*Epithelial Fine annotation:* After re-clustering, we obtained eight epithelial clusters. We identified C1 assigned as EMT epithelia with expression of collagen markers^125,126^, C3 with expression of vimentin (VIM) and SOX4^127,128^; C4 with ciliated epithelial markers PIFO+, and C7 with expression of MUC5B+/TFF3+ indicating progenitor cells^14^.

### Differential abundance, PCA composition, differential gene expression and pathway analysis

Differential abundance in each specific cell type was assessed using t-test and compare_means, provided by the package ggpubr (v 0.6.0) resulting in adjusted p-values per each menstrual phase or disease comparison.

PCA composition was obtained by calculating the ratio of each specific cell type compared to the corresponding broad cell type for every sample. This was done for stromal, epithelial and immune subsets.

Differential gene expression analyses were conducted using the FindMarkers function in Seurat with the method MAST^129^ (v. 1.18). The covariates associated with de-identified patient ID and library prep were included, and p-values were adjusted using the Benjamini-Hochberg method. DGE between endometriosis and pristine controls was performed in cell populations with two or more controls. Broad cell types and specific cell types were analyzed, using single-cell objects after filtering and sub-setting (see above) for immune, stromal fibroblast, VSM, ciliated and unciliated epithelial cells. Endothelia cells analysis was performed based on object containing all cells.

For pathway analysis, genes with an adjusted p-value less than 0.05 were selected. To calculate enrichment in GSEA we used clusterProfiler^130^, using “BP” as ontology, with pathways chosen based on a Benjamini-Hochberg adjusted p-value threshold less than 0.05.

### Cell communication analysis

In silico communication analysis was performed with CellChat^54^ (v1.6.1) using the CellChatDB.human as the database and populations as previously defined as broad non-immune (Stromal Fibroblast, VSM, Unicliated Epithelia, Ciliated Epithelia, and Endothelia) and broad immune. Each category (proliferative endometriosis, proliferative control, secretory endometriosis, and secretory control) was processed independently. Following standard procedures, we computed Communication Probability and filtered cell-cell communication with less than 10 cells. Cell objects were then merged and compared, stratifying by menstrual cycle. Data for visualization was retrieved by using the function subsetCommunication.

## Data Availability

Data available in CZ cellxgene (https://cellxgene.cziscience.com/ ). ScRNA-seq datasets to reproduce all figures, can be accessed and downloaded through the web portal.

## Code availability

The code for clustering, annotation, downstream analyses (including differential gene expression, differential abundance, and pathway enrichment), as well as generating figures, is available on GitHub https://github.com/almonteloya/Endometriosis_sc

## Acknowledgements

This work has been supported by NIH NICHD [P50 HD055764]. F.V. discloses support for the research of this work from Carlos III Institute of Health grant [PI21/00528 and PI24/01784], Spanish State Research Agency [CNS2022-135696], Conselleria de Innovacion, Universidad y Ciencia [CIAICO 2022/242], F.V and C.S disclose support from the European Commission Grant Agreement [101080219-2].

We thank INCLIVA Biobank (Hospital Clínico de Valencia) for providing subset of the samples used in this project.

The authors thank all members of the Sirota and Fragiadakis Lab for useful comments and feedback. We also thank Jessica Neely MD, Camilla Wibrand and Emily Teets for helpful advice.

## Author Contributions

Conceptualization by A.A, W.W, S.H, U.K, C.P, T.T.O, D.H, J.I, A.C, M.S, G.F, L.G. Data Curation A.A, S.H, W.W. Formal Analysis A.A, W.W, E.T. Funding Acquisition S.H, C.S, F.V, L.G. Sample Collection S.H, C.S, F.V, C.N. Sample Processing S.H, D.K, K.C.V, A.C. Supervision A.C, M.S, G.F, L.G. Validation X.T, E.T. Writing Original Draft, A.A, X.T, B.L. Manuscript Editing and Reviewing: All authors.

