## Supplementary material for "Single-Cell Profiling Reveals Altered Endometrial Cellular Features Across the Menstrual Cycle in Endometriosis Patients": Suplemental Notes and Figures

### Supplementary Information

#### Supplementary Notes

##### The transcriptional similarity between endometriosis and fibroids

Principal component analysis of cell composition showed separation across the menstrual phase (**Fig. 2D**), as well as separation between endometriosis, fibroids and pristine controls (**Fig. 2E**). To further investigate transcriptional similarities, we compared gene expression changes between secretory and proliferative samples across the three disease statuses (**SFig. 2A**). We first identified DEGs between the proliferative and secretory phases within each disease status **(STable 4**) (**SFig. 3**). Hierarchical clustering based on the significant DEGs showed greater similarity between fibroid controls and endometriosis in both stromal fibroblasts and unciliated epithelial cells, while pristine controls formed a separate cluster (**SFig. 2B,C**). Additionally, we found limited overlap in phase-related DEGs across disease groups, with most transcriptional changes being unique to a particular group (**SFig. 4).**

To assess the potential of fibroids as pathogenic controls, we performed differential gene expression analysis between fibroids and pristine controls **(STable 5)** (**SFig. 5A,B)** and compared these findings to DEGs in endometriosis. In the secretory phase, fibroids and endometriosis shared a general overlap in DEGs (**SFig.5C-G)**. Notably, in stromal fibroblasts, 54% of significant genes in endometriosis were also upregulated in fibroids (**SFig.5D**). Given the transcriptional similarities, we opted to use pristine controls for comparison with endometriosis cases.

#### Cellular gene expression changes specific to endometriosis severity

Endometriosis varies in severity based on the characteristics of lesions and adhesions; it is categorized into four stages: minimal (I), mild (II), moderate (III), and severe (IV). In the minimal and mild stages, women generally have superficial implants and a few adhesions. By contrast, those with moderate or severe endometriosis usually present with cysts and more pronounced adhesions.

To determine whether endometriosis severity is associated with distinct gene expression profiles, we stratified samples by menstrual cycle phase (proliferative vs. secretory) and compared controls to endometriosis samples of varying severity: mild (stage I/II), severe (stage III/IV), and all stages combined (**SFig. 8A**) (**STable 6,7).** Then, we quantified the similarity between these DEGs and those from all endometriosis samples by calculating the correlation of their log fold changes (**SFig. 8B-F**). In the proliferative phase, mild endometriosis samples showed a high Pearson correlation on average ($r >= 0.96$) across all non-immune cell types. However, in severe cases, the correlation dropped to an average r of 0.78, indicating that general endometriosis samples better represent the transcriptional changes in mild rather than severe endometriosis. In the secretory phase, we found a stronger correlation (average $r=0.96$) across both mild and severe cases, indicating that general endometriosis samples effectively captured severity-specific gene expression changes. Further analysis of severe endometriosis samples in the proliferative phase could be explored; however, based on these results, we decided to continue the analysis using all endometriosis samples rather than stratifying by severity, since it generally captures broad differences.

#### Supplementary Figure Legends

**Figure S1. Menstrual Cycle Phases and Disease Conditions Mapped via UMAP**

A. Expression of selected markers across major cell types
B. Global UMAP depicting menstrual phase density.
C. Global UMAP showing cell density of endometriosis, fibroids, and pristine control samples.
D. Cell distribution of endometriosis, fibroid controls, and pristine controls.
E. Cell distribution of endometriosis, divided by severity, fibroid, and pristine controls.
F. Cell distribution of the menstrual phase.
G. Cell distribution of libraries across cell groups.

**Figure S2. The effect of endometrial pathology on stromal fibroblasts and unciliated epithelial cells varies by cycle phase**

A. Study design for differential gene expression analysis. Samples were stratified by disease status (endometriosis, fibroid controls, and pristine controls) and compared between the secretory and proliferative phases. Differential expression was assessed with adj. p-val $<0.05$ and $|\text{log2FC}| > 0.25$. Differential expression analyses were performed using MAST.
B. Heatmap of upregulated genes between the secretory and proliferative phases, stratified by disease status endometriosis, fibroid controls, and pristine controls). Genes included were present in at least two groups. Hierarchical clustering displays gene expression patterns along with disease status, showing the stromal fibroblast population.
C. Heatmap showing unciliated epithelial cells, highlighting the upregulated genes between the secretory and proliferative phases, stratified by disease status.

**Figure S3. Menstrual cycle phase-specific differential gene expression in non-immune broad cell types**

A-O. Samples were stratified by disease status (endometriosis, fibroid controls, and pristine controls) and compared between secretory and proliferative phases. Differential expression was assessed with adj. p-val< 0.05 and |log2FC| > 0.25. Differential expression analyses were performed using MAST.

**Figure S4. Cycle phase-specific genes are disease status-specific.**

A-E. Samples were stratified by disease status (endometriosis, fibroids, and pristine controls) and compared between secretory and proliferative phases. Showing the overlap of signal per cell type.

**Figure S5: Fibroid gene expression shows differences with pristine controls across menstrual phases.**

A. Study design for differential gene expression analysis. Samples were stratified by menstrual status and compared between fibroid controls and pristine controls. Differential expression was assessed with adj. p-val<0.05 and |log2FC| > 0.25. Differential expression analyses were performed using MAST

B. Number of upregulated and downregulated genes in each broad cell type.

C-G. Overlap between upregulated and downregulated genes stratified by menstrual phase, comparing [fibroid controls vs. pristine controls] and [endometriosis vs. pristine controls] per cell type.

**Figure S6: Endometriosis vs pristine controls in broad non-immune cell types stratified by menstrual phase**

A-J. Differential gene expression analysis. Samples were stratified by menstrual phase to compare pristine controls with endometriosis samples. Differential expression was assessed with adj. p-val<0.05 and |log2FC| > 0.25. Analyses were performed using MAST. Showing each broad cell type.

**Figure S7: Overlap by phase DEG of endometriosis vs pristine controls in broad non-immune cell types**

A-E. Overlap of DEG genes between menstrual phases. Comparing pristine controls with endometriosis samples.

F. Endothelia upregulated pathways in endometriosis in the proliferative and secretory phases. Using GSEA, adj. p-val< 0.05

**Figure S8: Stratifying endometriosis cases based on severity correlates with previous results.**

A. Study design for differential gene expression analysis. Samples were stratified by menstrual phase. Differential expression was assessed with adj. p-val<0.05 and |log2FC| > 0.25. Differential expression analyses were performed using MAST.

B. Comparisons between 1) all endometriosis samples vs pristine controls, 2) Mild endometriosis vs pristine controls, and 3) Severe endometriosis vs pristine controls. C

C-F. Comparisons between the log fold change of using all endometriosis samples and stratifying by endometriosis severity. Showing broad cell types.

**Figure S9. Cluster composition and abundance of stromal fibroblast cells**

A. Heatmap showing significant genes in endometriosis, adj. p-val<0.05 and |log2FC| > 0.25, stratified by cycle phase. Showing genes intersecting across the menstrual phase.
B. Bar plots depicting the proportion of cells within stromal subclusters from samples in either the secretory or proliferative phase of the menstrual cycle.
C. Bar plots depicting the proportion of cells within stromal subclusters classified according to endometriosis and pristine controls.

D. Bar plots depicting the proportion of cells within stromal subclusters classified according to endometriosis severities or pristine controls
E. Bar plots depicting the proportion of cells within stromal subclusters classified by sample.
F. Bar plots depicting the proportion of cells within stromal subclusters classified by library.
G. Box plots showing the proportion of cells within each subpopulation of stromal cells. Stratified by menstrual phase. Comparisons were made using a T-test to assess proportional differences between endometriosis and pristine controls.
H. Stratified among control patients and endometriosis, showing changes between secretory and proliferative samples.

**Figure S10. Characterizing the Stromal subpopulation in the secretory phase**

A-H. Differential gene expression analysis across stromal subpopulations, stratified by the secretory phase, to compare endometriosis samples with pristine controls. Differential expression was evaluated with a significance threshold of adj. p-val< 0.05 and a |log2FC| > 0.25. Analyses were conducted using MAST.

**Figure S11. Characterizing Stromal Fibroblast subpopulation in the proliferative phase**

A-H. Differential gene expression analysis across stromal subpopulations, stratified by the proliferative phase, to compare endometriosis samples with pristine controls. Differential expression was evaluated with a significance threshold of adj. p-val< 0.05 and a |log2FC| > 0.25. Analyses were conducted using MAST.

**Figure S12. Cluster composition and differential abundance of epithelial cells**

A. Heatmap showing significant genes in endometriosis, adj. p-val<0.05 and |log2FC|> 0.25, stratified by cycle phase. Showing genes intersecting across the menstrual phase.

B. Bar plots depicting the proportion of cells within epithelial subclusters from samples in either the secretory or proliferative phase of the menstrual cycle.

C. Bar plots depicting the proportion of cells within epithelial subclusters classified according to endometriosis and pristine controls.

D. Bar plots depicting the proportion of cells within epithelial subclusters classified according to endometriosis severities or pristine controls

E. Bar plots depicting the proportion of cells within epithelial subclusters classified by sample.

F. Bar plots depicting the proportion of cells within epithelial subclusters classified by library.

G. Box plots showing the proportion of cells within each subpopulation of epithelial cells. Stratified by menstrual phase. Comparisons were made using a T-test to assess proportional differences between endometriosis and pristine controls.

H. Stratified among control patients and endometriosis, showing changes between secretory and proliferative samples.

**Figure S13. Characterizing the Epithelial subpopulation in the secretory phase**

A-E. Differential gene expression analysis across epithelial subpopulations, stratified by the secretory phase, to compare endometriosis samples with pristine controls. Differential expression was evaluated with a significance threshold of adj. p-val< 0.05 and a |log2FC| > 0.25. Analyses were conducted using MAST.

**Figure S14. Characterizing Epithelial subpopulation in the proliferative phase**

A,B. Differential gene expression analysis across epithelial subpopulations, stratified by the proliferative phase, to compare endometriosis samples with pristine controls. Differential expression was evaluated with a significance threshold of adj. p-val < 0.05 and a |log2FC| > 0.25. Analyses were conducted using MAST.

**Figure S15. Immune cells according to composition and markers.**

A. Bar plots depicting the proportion of cells within immune subclusters from samples in either the secretory or proliferative phase of the menstrual cycle.
B. Bar plots depicting the proportion of cells within immune subclusters classified according to endometriosis and pristine controls.
C. Bar plots depicting the proportion of cells within immune subclusters classified according to endometriosis severities or pristine controls
D. Bar plots depicting the proportion of cells within immune subclusters classified by sample.
E. Bar plots depicting the proportion of cells within immune subclusters classified by library.
F. Box plots showing the proportion of cells within each subpopulation of immune cells. Stratified by menstrual phase. Comparisons were made using a T-test to assess proportional differences between endometriosis and pristine controls. "ns" not significant.
G. Stratified among control patients and endometriosis, showing changes between secretory and proliferative samples.

**Figure S16. Ligand-receptor pair expression with innate lymphoid as sender**

A. Bubble plot of the probability of ligand-receptor interaction with innate lymphoid as a sender in the proliferative phase. TNF pathway highlighted.

### Supplementary Figures

#### Figure S1: Menstrual Cycle Phases and Disease Conditions Mapped via UMAP

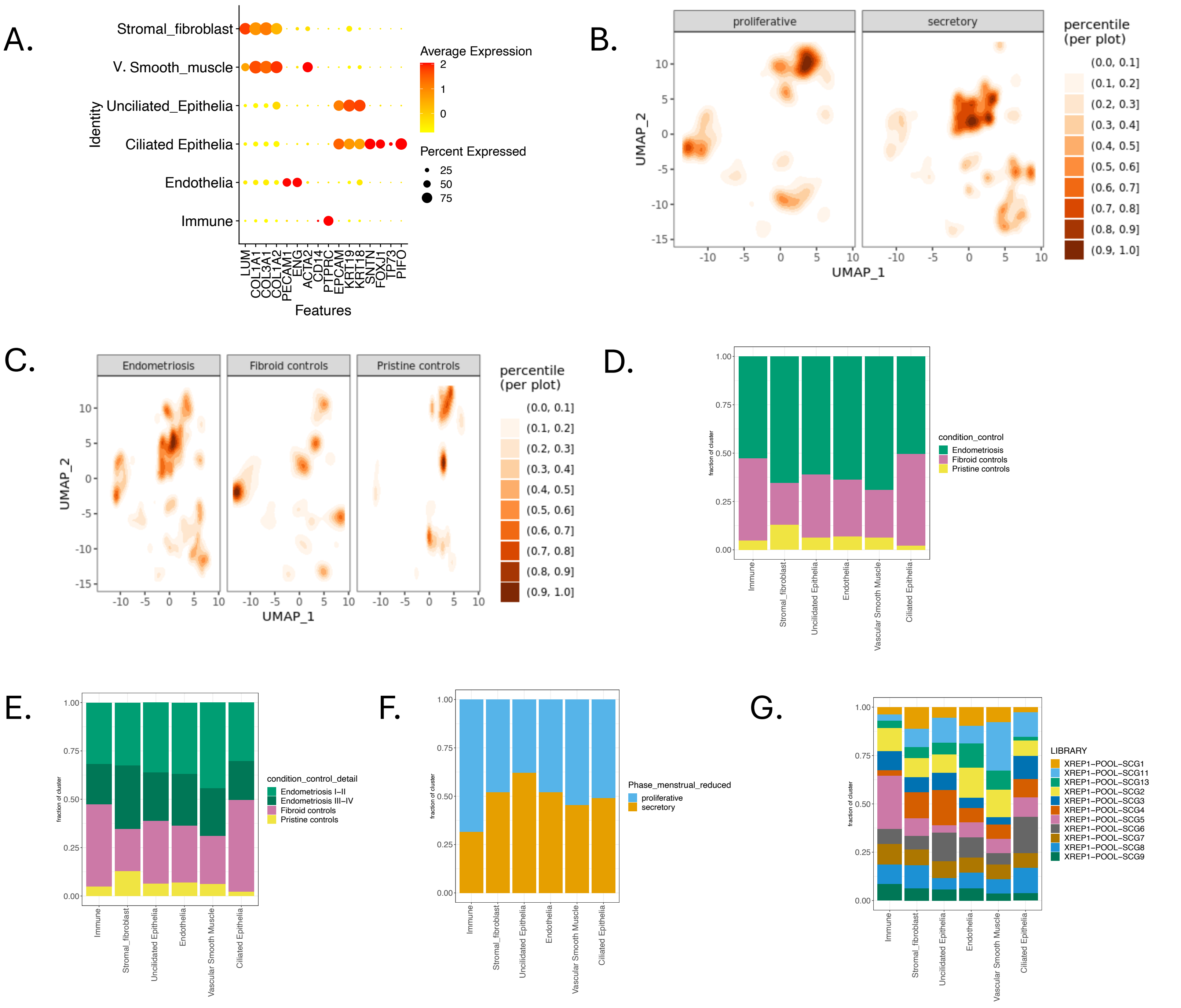

### Figure S2: The effect of endometrial pathology on stromal fibroblasts and unciliated epithelial cells varies by cycle phase

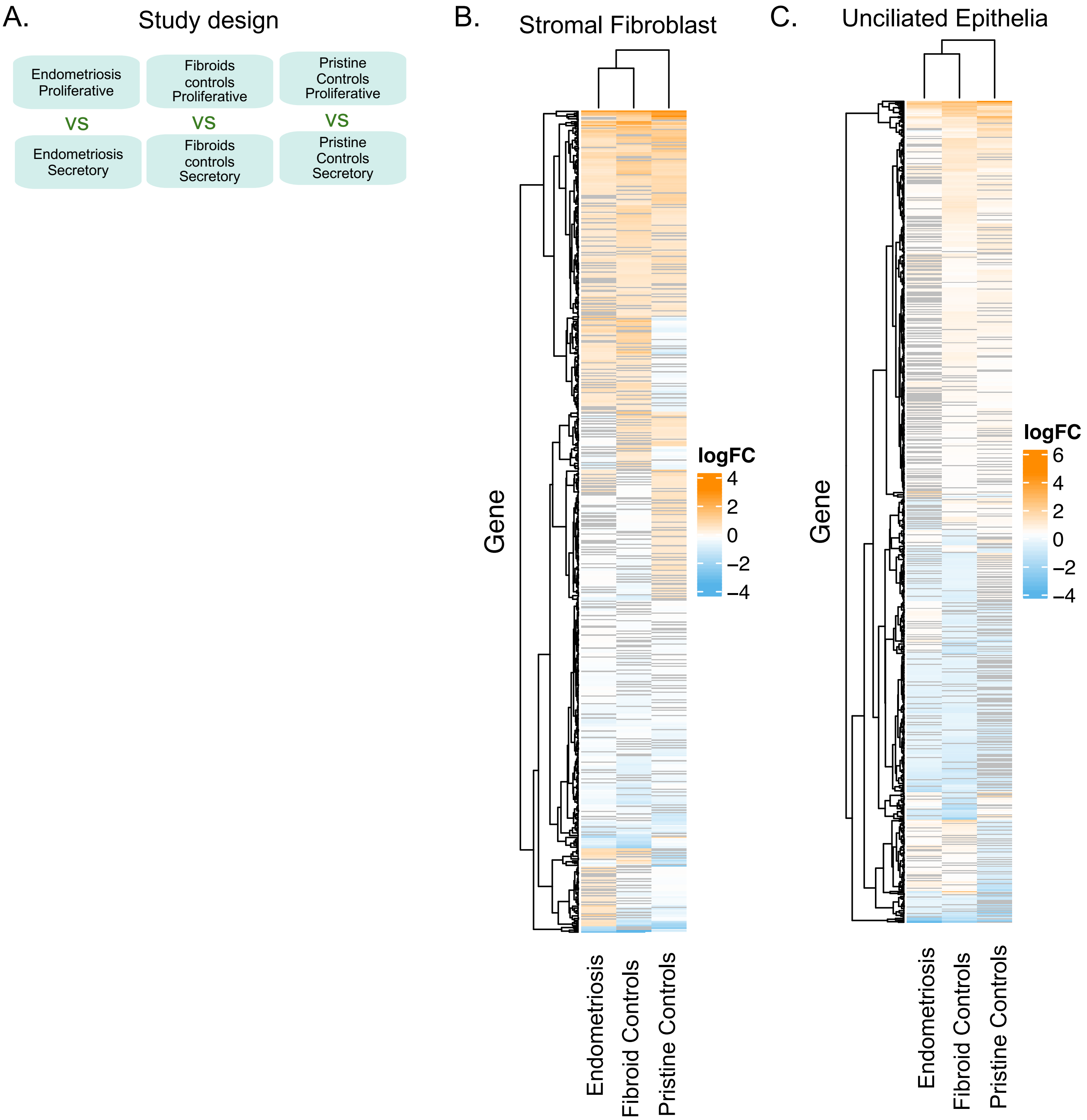

### Figure S3. Menstrual cycle phase-specific differential gene expression in non-immune broad cell types

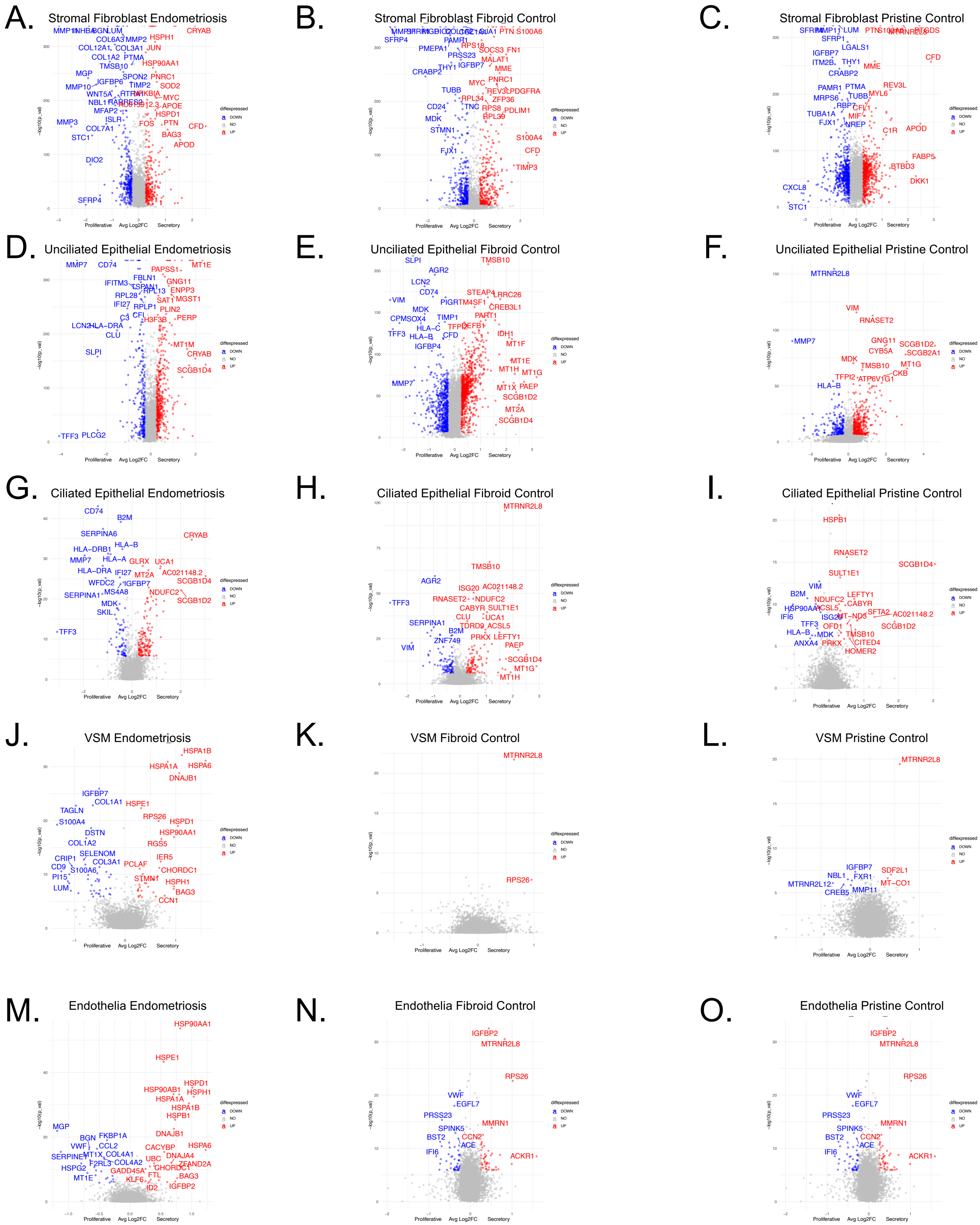

### Figure S4. Cycle phase-specific genes are disease status-specific.

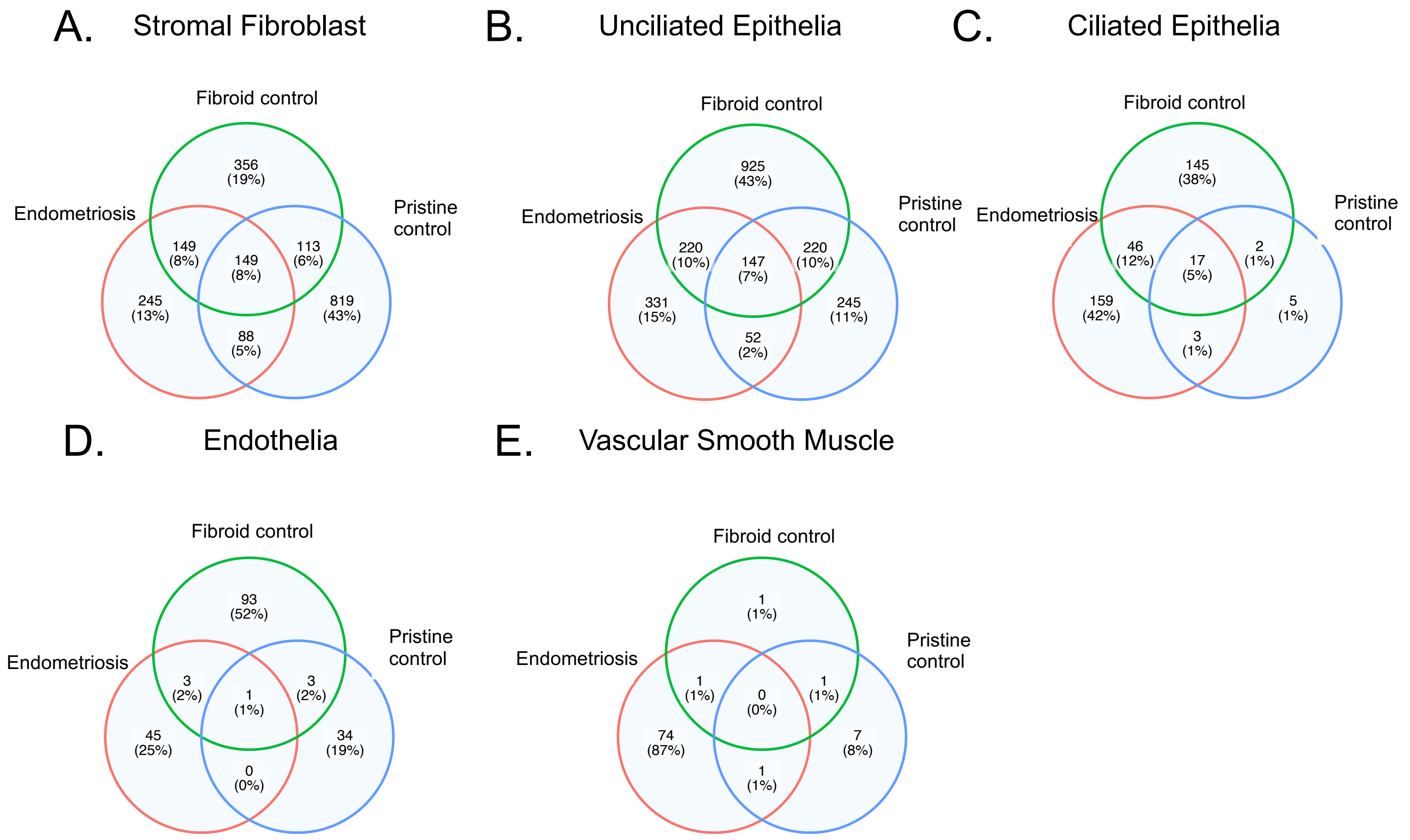

### Figure S5: Fibroid gene expression shows differences with pristine controls across menstrual phases.

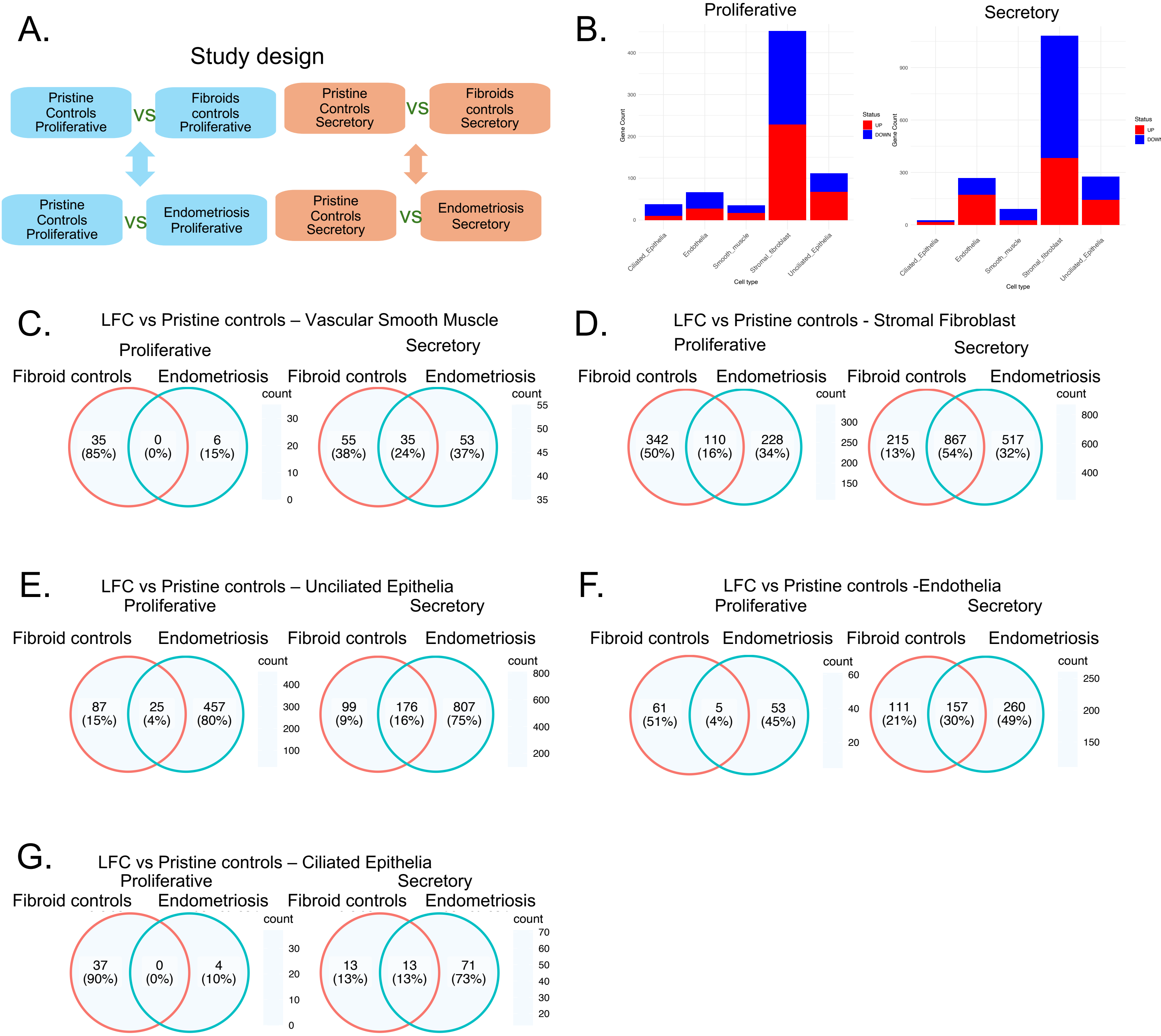

### Figure S6: Endometriosis vs pristine controls in broad non-immune cell types stratified by menstrual phase

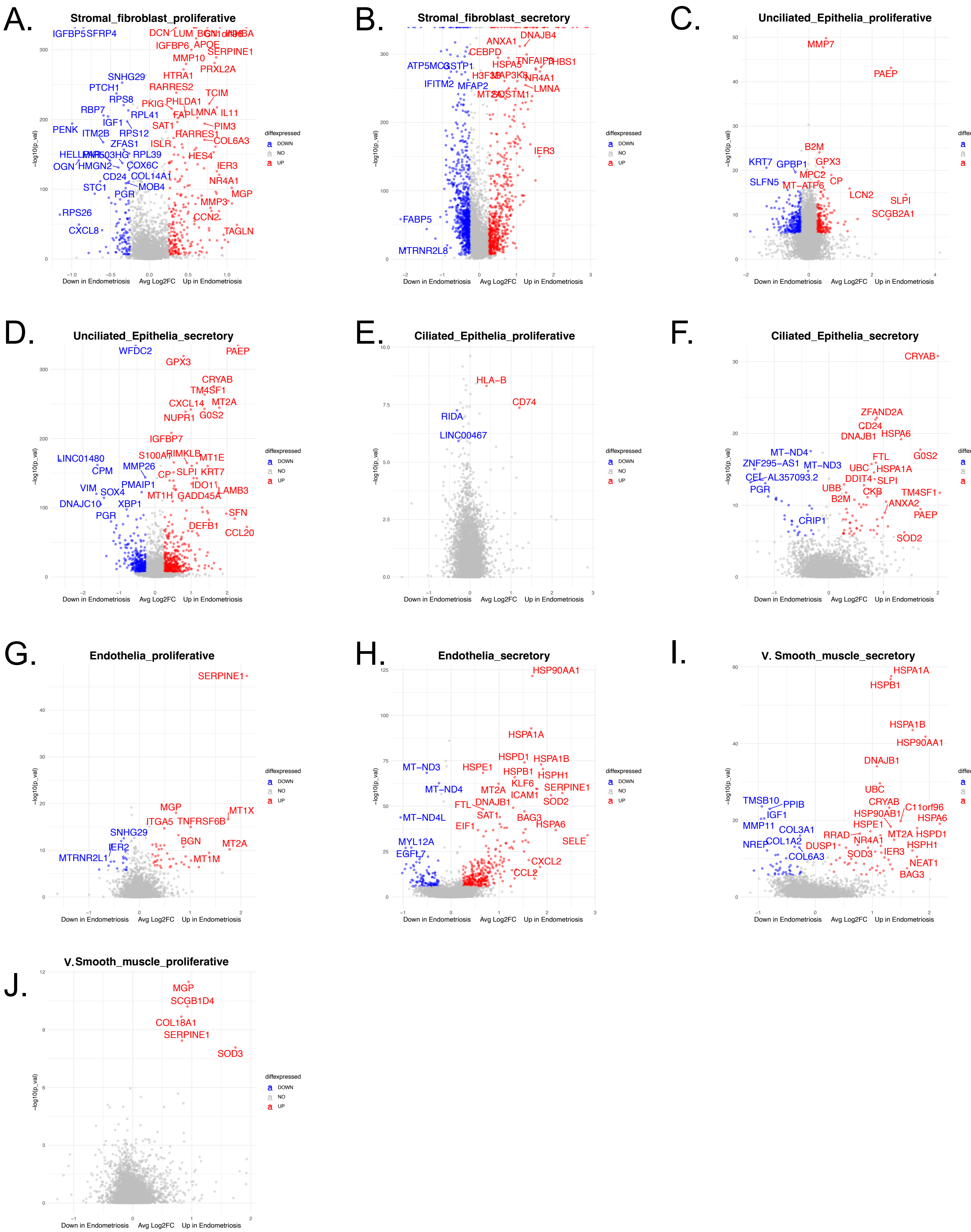

### Figure S7: Overlap by phase DEG of endometriosis vs pristine controls in broad non-immune cell types

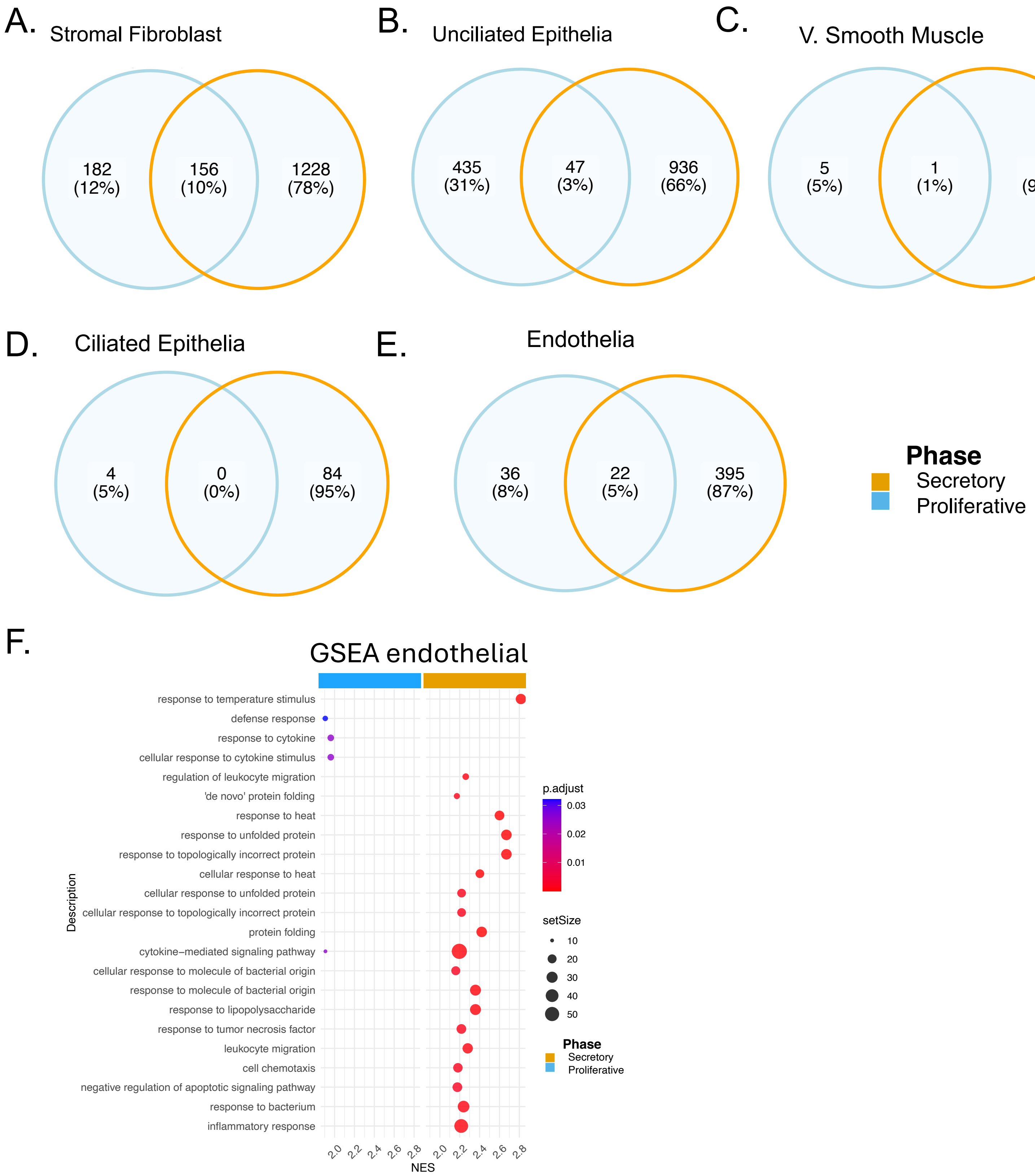

### Figure S8: Stratifying endometriosis cases based on severity correlates with previous results.

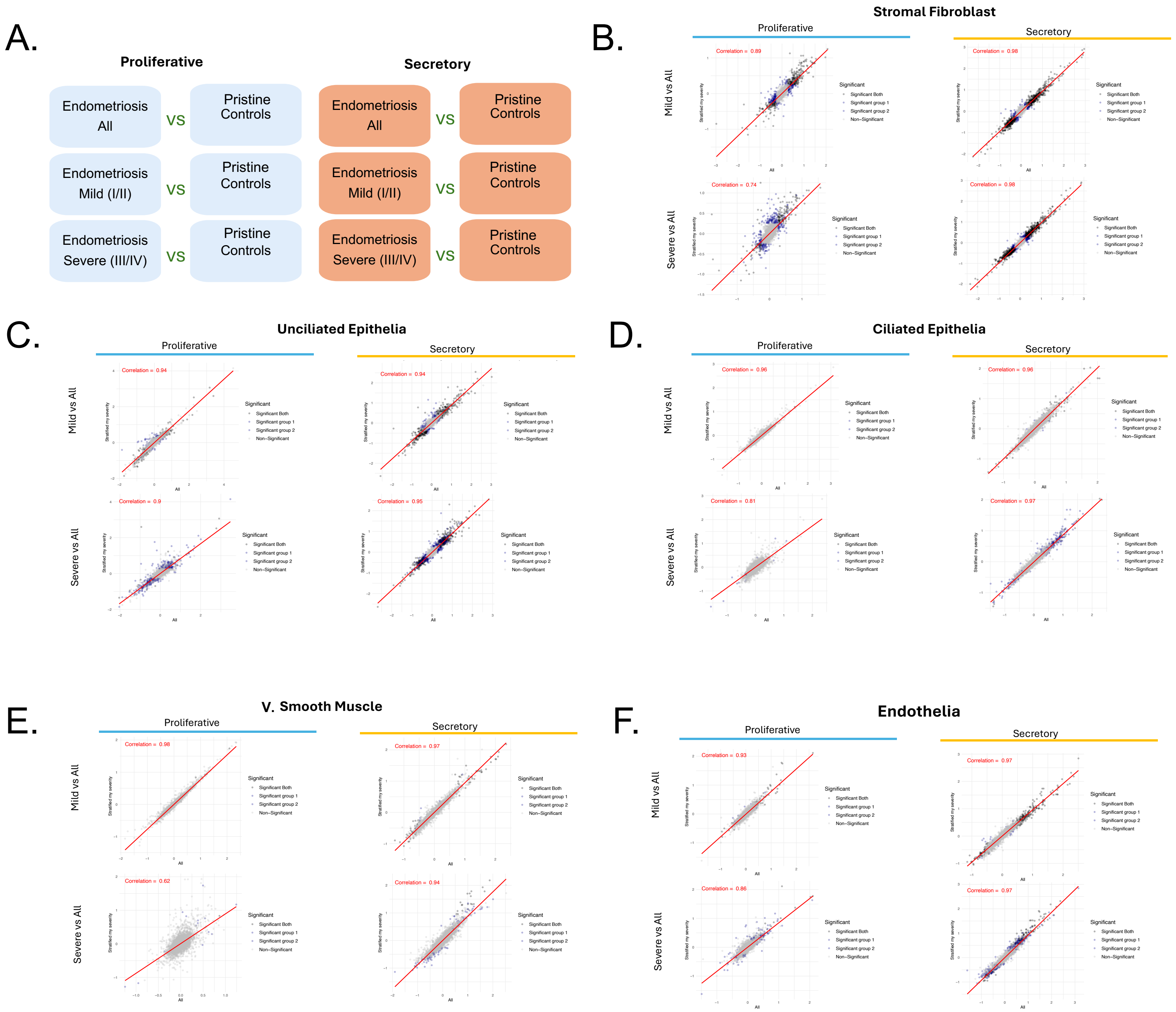

### Figure S9. Cluster composition and abundance of stromal fibroblast cells

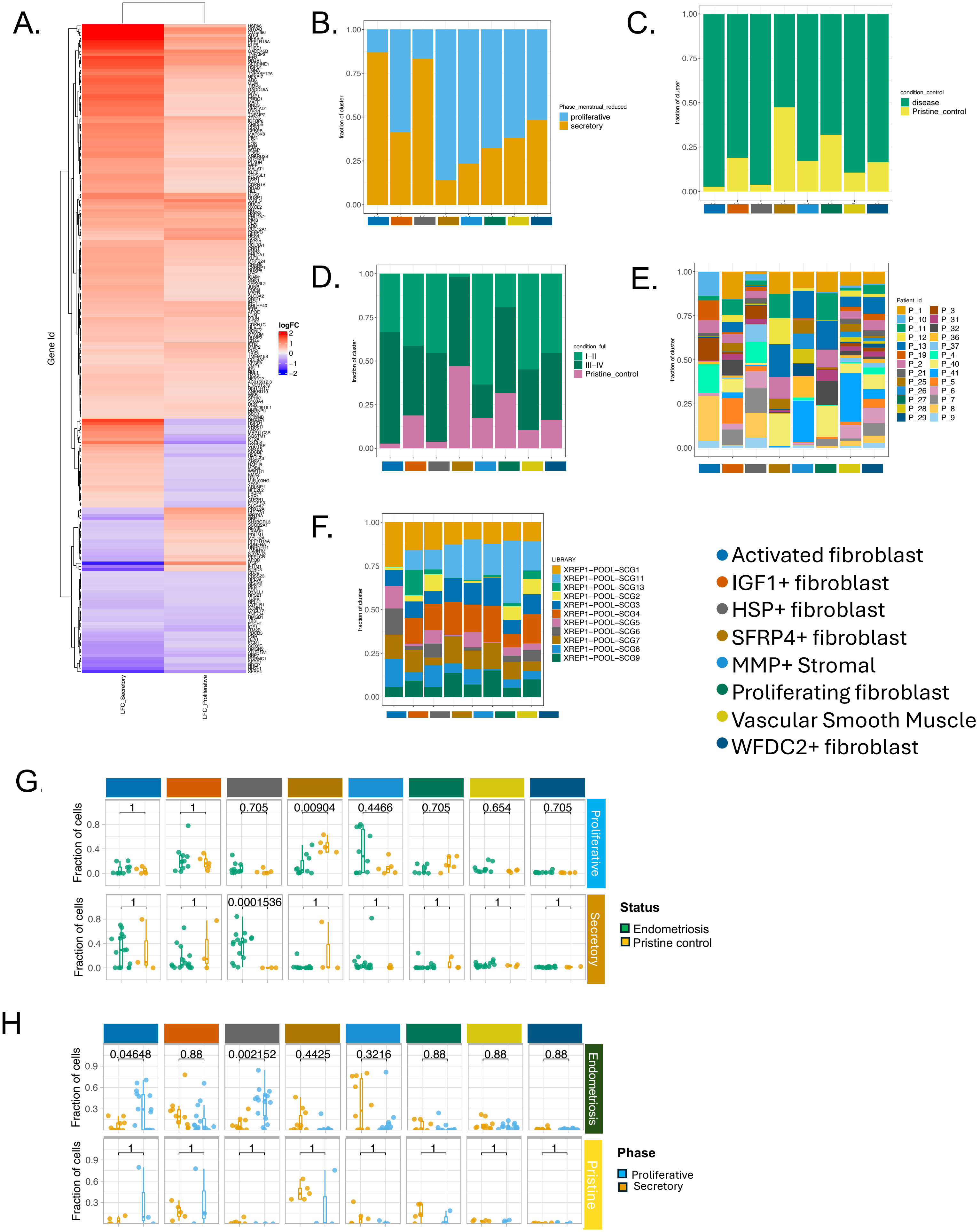

### Figure S10. Characterizing Stromal subpopulation in the secretory phase

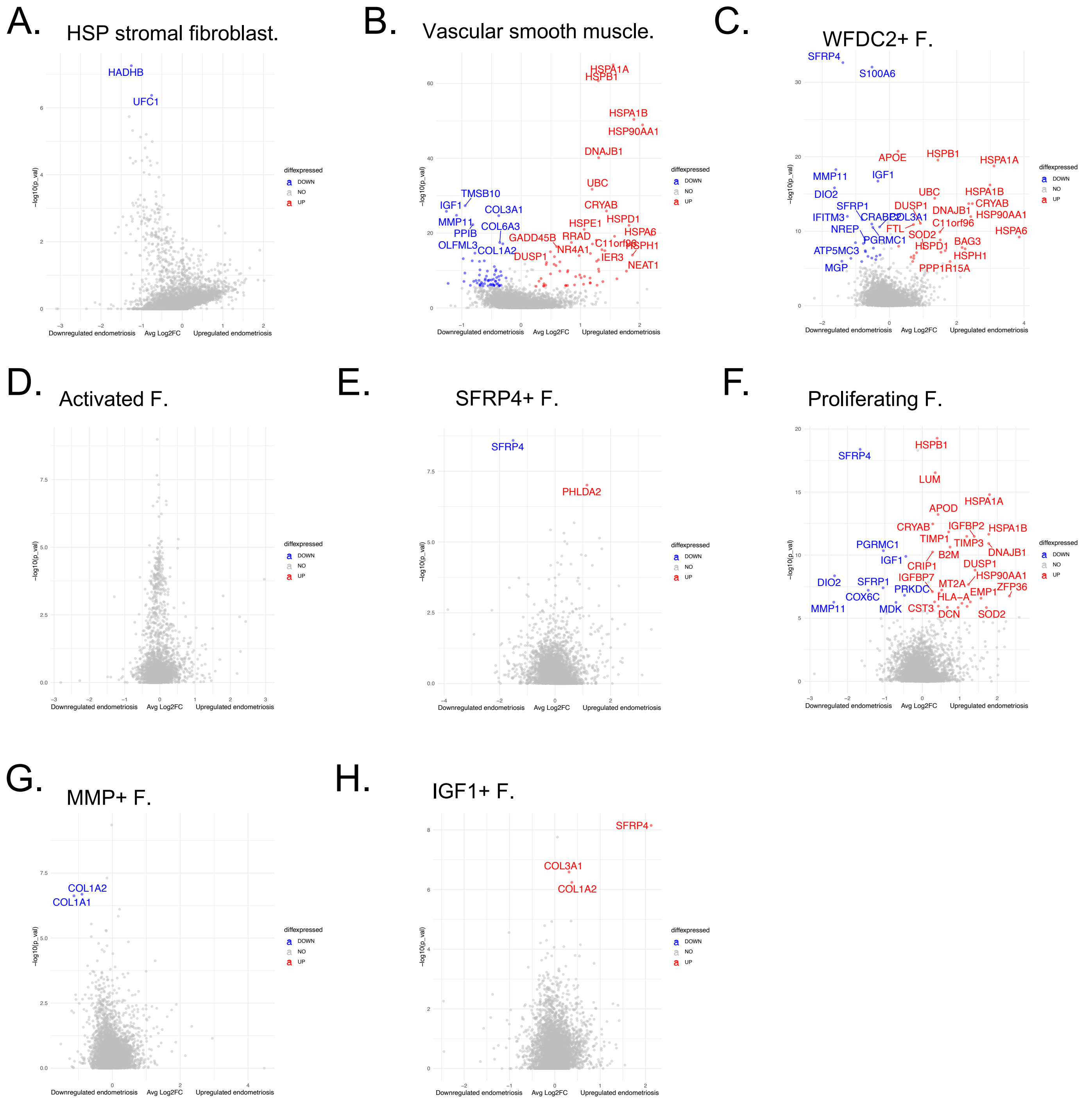

Figure S11. Characterizing Stromal Fibroblast subpopulation in the proliferative phase

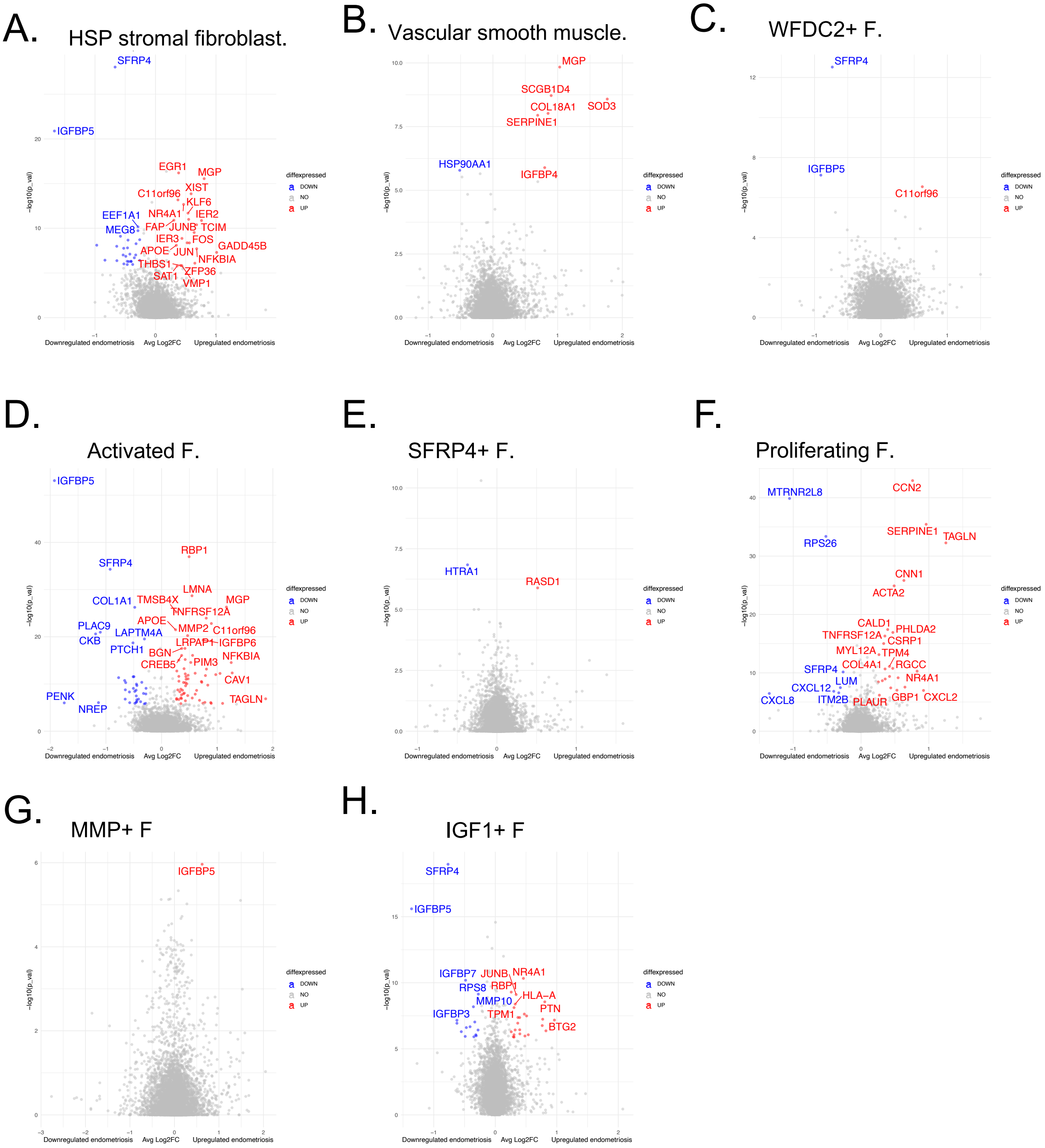

### Figure S12. Cluster composition and differential abundance of epithelial cells

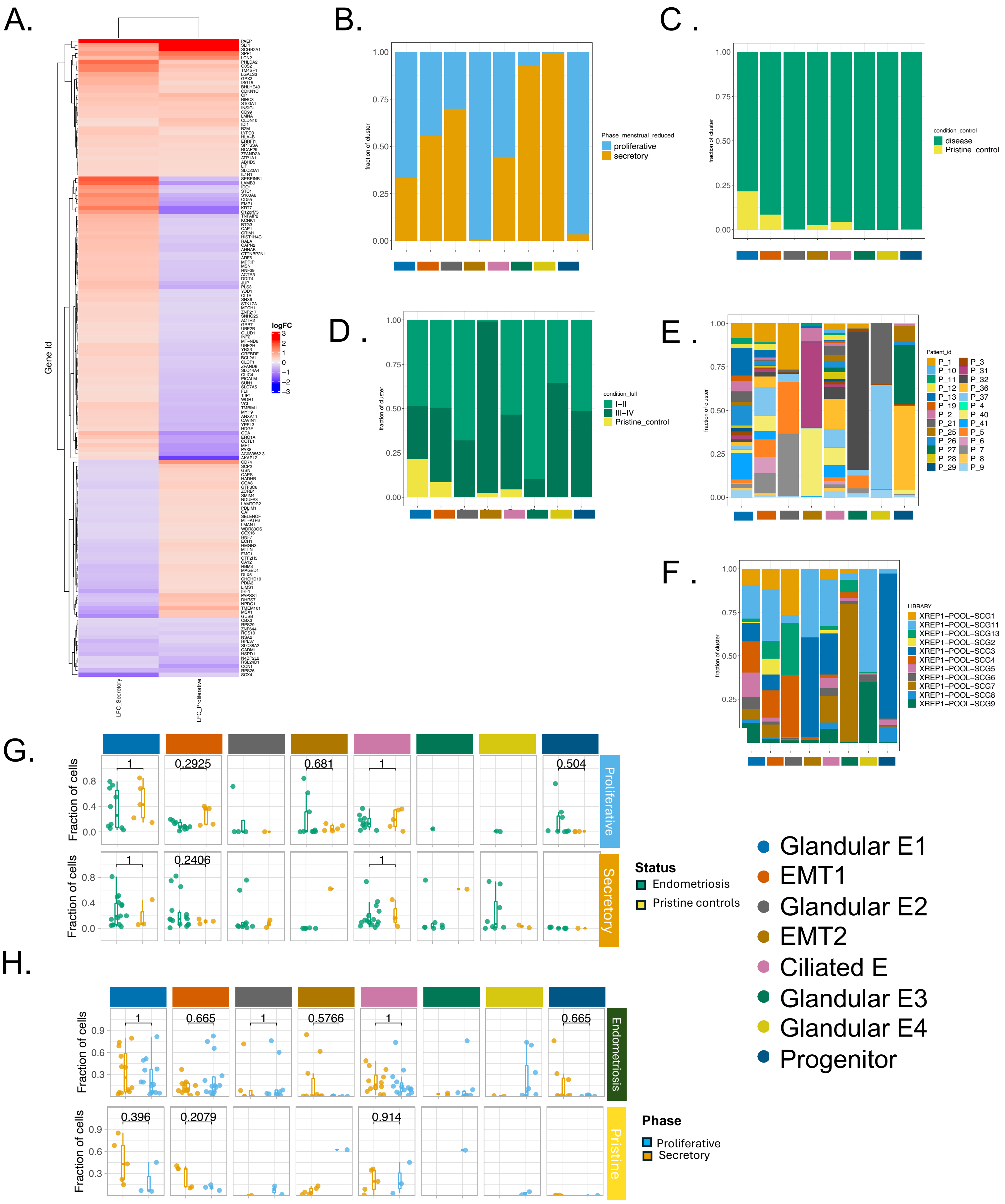

### Figure S13. Characterizing Epithelial subpopulation in the secretory phase

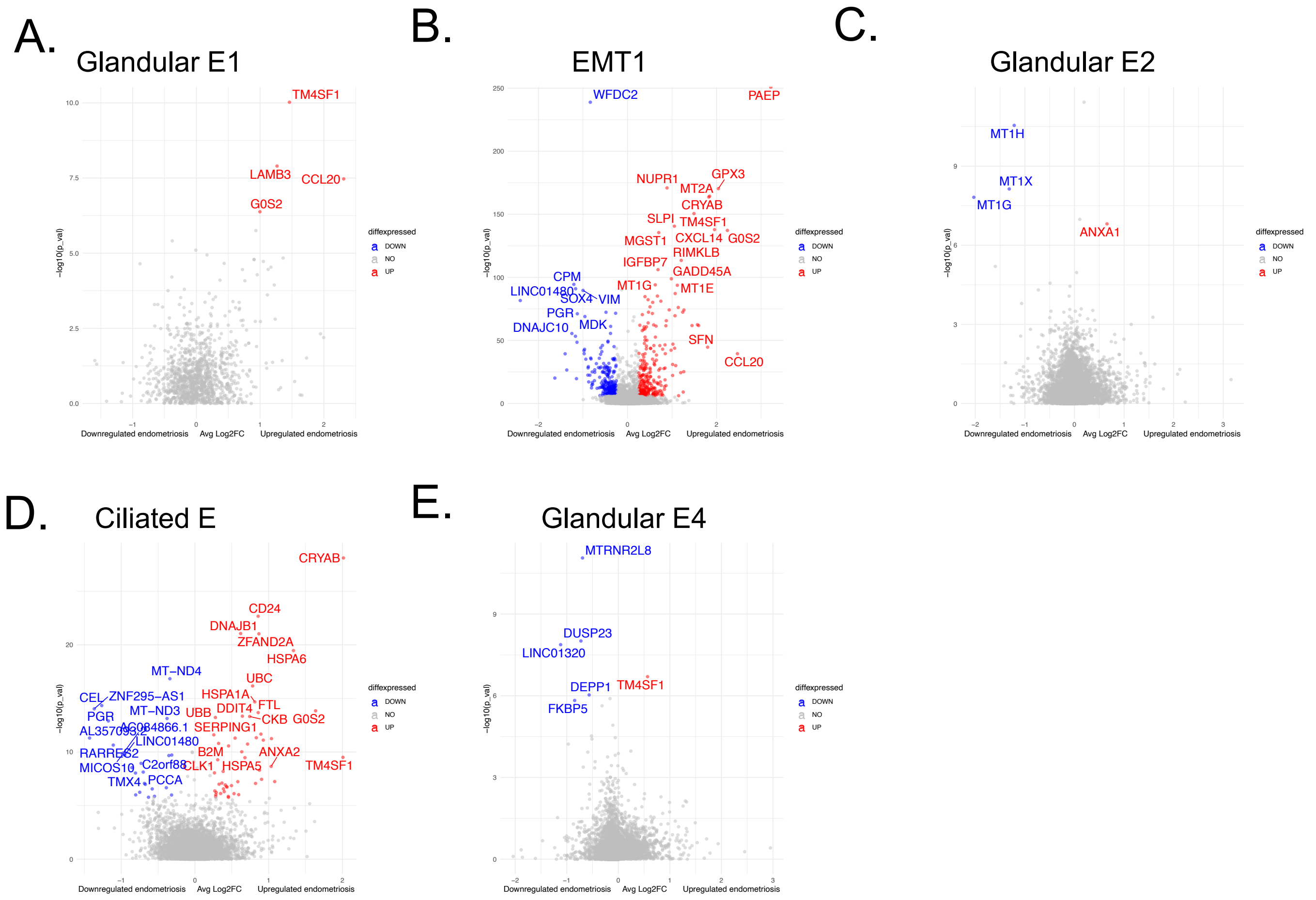

### Figure S14. Characterizing Epithelial subpopulation in the proliferative phase

A. B.

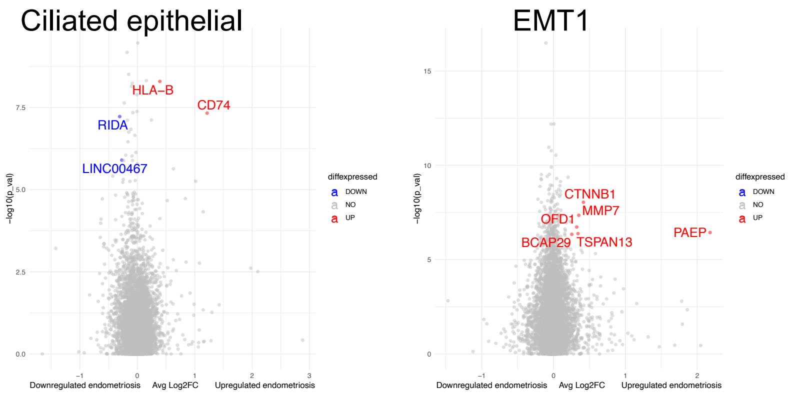

### Figure S15. Immune cells according composition and markers.

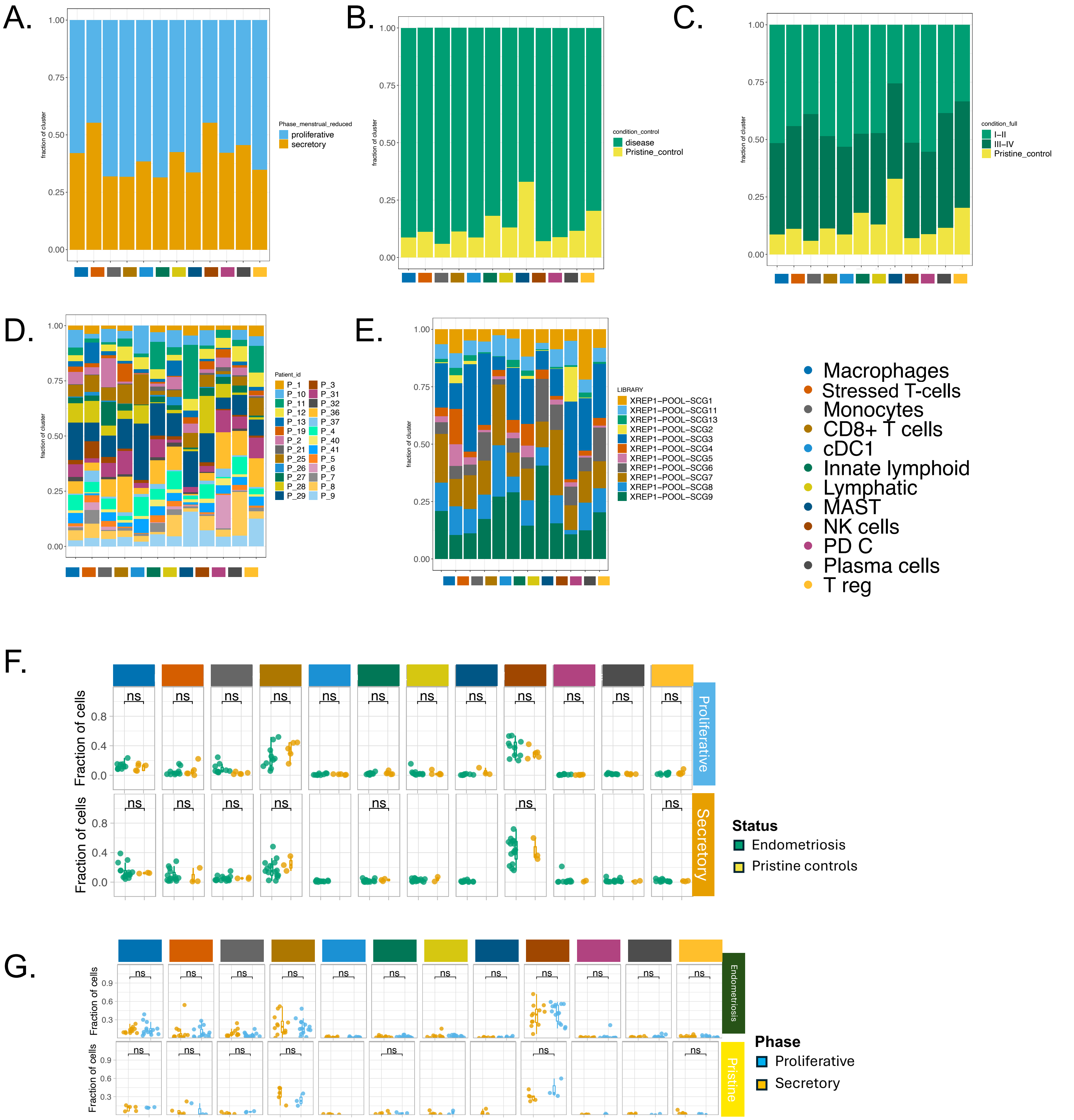

### Figure S16. Ligand-receptor pair expression with innate lymphoid as sender

A.

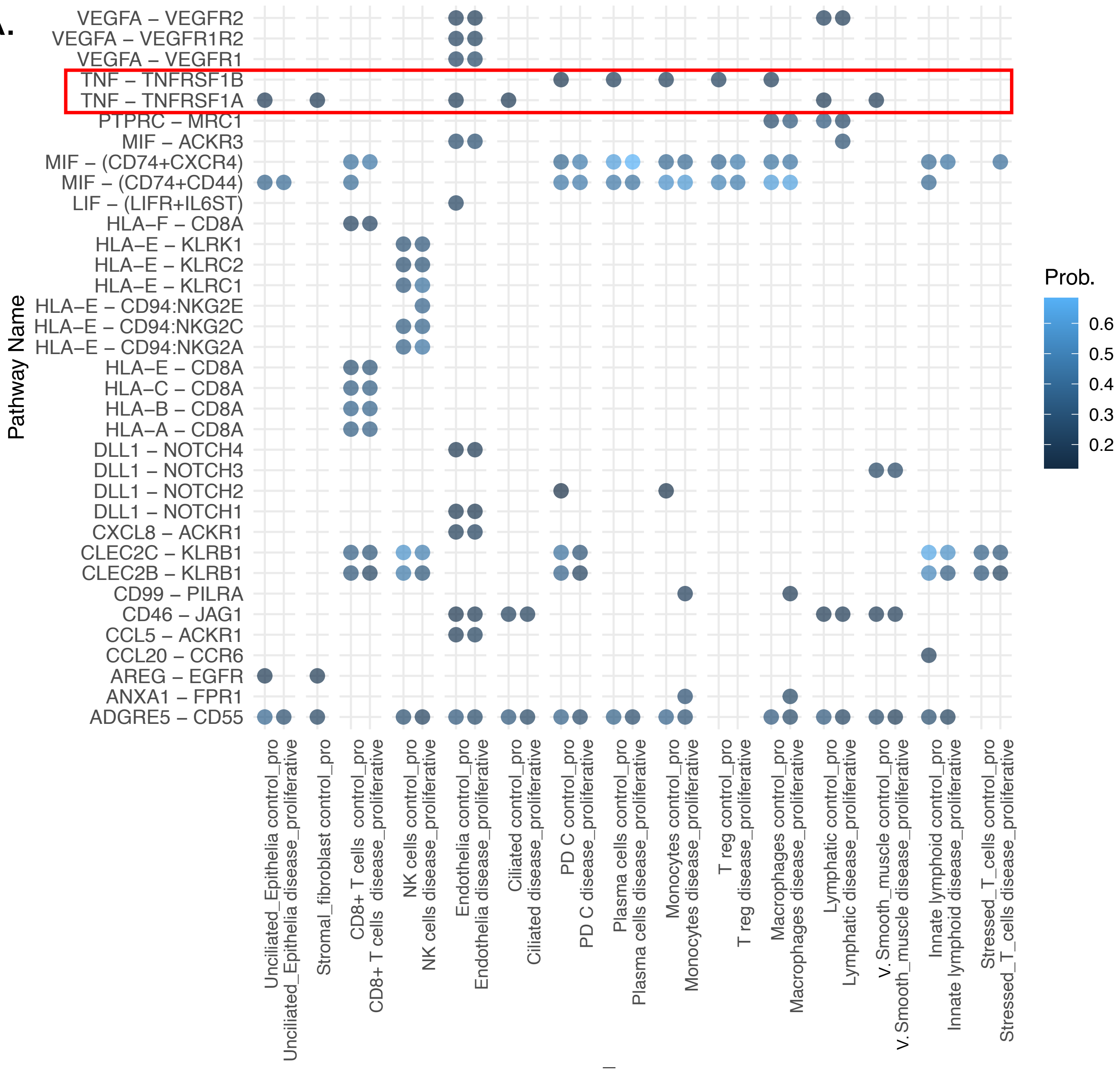
